# Chronic lysosome damage boosts interferon responses to Palbociclib dependent upon the mitochondrion

**DOI:** 10.1101/2025.03.13.642980

**Authors:** Mihaela Bozic, Tian En Lim, Carla Salomó-Coll, Arianna Karageorgiou, Alain J. Kemp, Katie Winnington-Ingram, Laura Murphy, Ashish Dhir, Ann Wheeler, Natalia Jimenez-Moreno, Simon Wilkinson

## Abstract

Acute lysosome damage triggers the endolysosome damage response (ELDR) in order to co-ordinate vesicle repair or removal by autophagy (lysophagy). However, it is unclear whether persistent damage, as occurs after chronic challenge to lysosome integrity, triggers wider cellular responses. Here, we show that longitudinal treatment with a lysosomotropic cancer therapeutic, the CDK4/6 inhibitor Palbociclib, invokes chronic lysosome damage in breast and lung cancer cells. Autophagy ameliorates but does not avert this phenotype, which persists over days. Damaged lysosomes form contacts with mitochondria, which correlates with mitochondrial stress and cytosolic efflux of immunostimulatory mitochondrial nucleic acids. Importantly, mitochondrial nucleic acid release is necessary for the anti-cancer interferon response to Palbociclib. In conclusion, chronic lysosome damage rewires cellular signalling responses in a mitochondrion-dependent manner and this effect should be considered when assessing the cellular actions of cancer therapeutics.

**Summary statement:** Bozic *et al* suggest that lysosome damage can trigger interferon responses dependent upon mitochondrial release of immunogenic nucleic acid. This is associated with damaged lysosome-mitochondrion contacts and is prevented by autophagy.

## Introduction

Lysosomes are single membrane-bound hydrolytic organelles that act at the termini of endocytic and autophagic pathways ^1^. Although rupture of pathogen-containing phagolysosomes is a feature of some infections ^2^, it is also well established that, in a sterile context, widespread loss of integrity of lysosomes can result in toxic leakage of hydrolytic proteases to the cytoplasm and subsequent cell death ^3^. However, acute, sublethal loss of lysosome membrane integrity typically induces the endolysosome damage response (ELDR), a collective term for pathways facilitating membrane stabilisation and repair ^4–6^, new lysosome biogenesis ^7^ or lysophagy ^8^, the latter being the process of selective degradation of the damaged lysosome via autophagy ^9^. Lysophagy requires evolutionarily-conserved core autophagy proteins that facilitate generation of nascent autophagosome vesicles and the concurrent sequestration of damaged lysosome cargo therein ^10,11^. Mature autophagosomes fuse with healthy lysosomes to facilitate degradation of sequestered cargo. Damaged lysosomes are initially recognised for degradation via “eat-me” signals ^8,12^. These include recruitment of galectin proteins, for instance Galectin 8 (GAL8) ^2,13^, to glycosylated lysosomal membrane proteins, and conjugation of polyubiquitin chains to lysosomal proteins ^14,15^.

Mechanistically, acute damage to lysosomes leads to galectin- and polyubiquitin-mediated recruitment of molecules that mediate lysophagy ^2,7,13,16–21^, in particular adaptors that bind directly to ATG8 (LC3/GABARAP) proteins conjugated to nascent autophagosome membranes ^22^ and which also recruit additional factors upstream of ATG8 conjugation ^23–26^. Indeed, deletion of *ATG5* (Autophagy Related 5), which prevents ATG8 conjugation to autophagosomal (and lysosomal) membranes, is effective in stabilising damaged lysosomes by ablating lysophagy and also by compromising ATG8-mediated recruitment of repair factors ^27,28^. Importantly, the removal and repair responses described above are engaged acutely, within minutes of lysosomal damage.

Notably, while extensive lysosome damage provokes cell death, linked to neurodegeneration ^1,3^ and disorders in other tissues such as kidney ^9^, it is known that lysosome damage *per se* is not necessarily lethal to cells ^29^, and the broader cellular consequences of chronic — yet sublethal — damage are unclear. This is particularly true in the context where the acute restorative ELDR functions, such as those dependent upon *ATG5*, are overwhelmed. Recent data indicate that programmed signalling responses to lysosome damage may exist ^13^. For instance, lysosome rupture triggers RNA stress granule formation via Protein Kinase R (PKR) activation, which may maintain cell survival, over at least the first four hours of lysosomal challenge ^30,31^. However, on the whole, mechanistic studies have focussed heavily on acute damage after treatment with tool compounds, such as the membrane lytic peptide L-Leucyl-L-Leucine methyl ester (LLOMe) ^9^. Importantly, numerous cancer therapeutics accumulate within lysosomes in either the cancer cells or other cell types. Generally, this phenomenon is viewed as detracting from therapeutic output by sequestering molecules from their primary targets. Nonetheless, it is possible that this may be consequential for the mechanism of the drugs. For instance, exposure of primary myeloid cells with lysosomotropic tyrosine kinase inhibitors may trigger lysosome damage and inflammasome activation, potentially contributing to treatment efficacy ^32^.

Palbociclib (Pb) is an important anti-cancer drug that is sequestered from its enzymatic target, the cell-cycle progression kinase CDK4/6 (cyclin-dependent kinase 4/6), due to lysosomal trapping ^33^. Strikingly, in addition to cell-cycle inhibition, Palbociclib also engages an anti-tumoural interferon response ^34–36^. Signalling occurs via detection of immunostimulatory moieties, purportedly from nuclear-derived double-stranded (ds) RNA ^37^, by cytosolic pattern recognition receptors (PRRs) ^38^, including RIG-I (retinoic-acid inducible gene 1) working in tandem with MAVS (mitochondrial antiviral signaling protein) ^39–42^. This action of Palbociclib has been shown to drive anti-tumour immunity ^34^. Indeed, discovering triggers and mechanisms of sterile interferon responses via PRRs is an important research goal in cancer therapy ^43,44^. Other potential triggers could include exogenous DNA, nuclear DNA, and mitochondrial DNA or dsRNA ^45–48^.

Here, we reveal a link between chronic lysosome damage and interferon signalling. We show that chronic lysosome damage can be provoked by Palbociclib, independently of CDK4/6, as well as the tool compound LLOMe. Chronic lysosome damage results in persistent interorganellar contacts between ruptured lysosomes and mitochondria. Lysosome damage is associated with stress responses within mitochondria and, ultimately, accumulation of immunostimulatory mitochondrial nucleic acid in the cytosol, which we show is a requisite event for productive interferon responses. However, in the case of Palbociclib the interferon responses are synergistically facilitated by concomitant CDK4/6 inhibition. Overall, our data show that mitochondria act as priming hubs in triggering of sterile interferon responses associated with chronic lysosome damage. This molecular mechanism has implications for understanding of — and for amelioration or enhancement of — the cellular response to various chronic insults, including cancer therapeutics.

## Results

### Palbociclib induces chronic lysosome damage and ELDR

We hypothesised that Palbociclib (Pb) might trigger lysophagy since it accumulates in lysosomes and causes swelling ^33^, i.e., potentially indicative of lysosome damage. Importantly, cancer treatments are applied chronically, over days and weeks, rather than on the acute minute-to-hour scales typically employed in lysophagy studies. We thus examined engagement of lysophagy flux by 24 h treatment with Pb, alongside the established lysosome-damaging compound LLOMe; these assays were performed using a new, sensitive reporter consisting of GAL8 fused to the C-terminus of SRAI (signal-retaining autophagy indicator, Fig. 1A) ^49,50^. Similar to LLOMe, Pb treatment was discovered to trigger lysophagy flux in A549 lung and MCF7 breast cancer cells (Figs. 1B, C, S1). Corroborating this, we also tested GAL8 fused to the conventional mRFP-GFP autophagy flux probe, and enumerated discrete individual autolysosomes containing damaged lysosome remnants (mRFP^+^ GFP^-^ galectolysosomes, wherein the acidic lysosome environment quenches the GFP signal, Fig. 1D, E). Again, these were induced by Pb treatment; additionally, consistent with elevated lysophagy flux, they were dependent upon *ATG5* (non-targeted wild-type, NT, compared with Δ*ATG5* cells, Fig. 1D, E).

**Figure 1.**
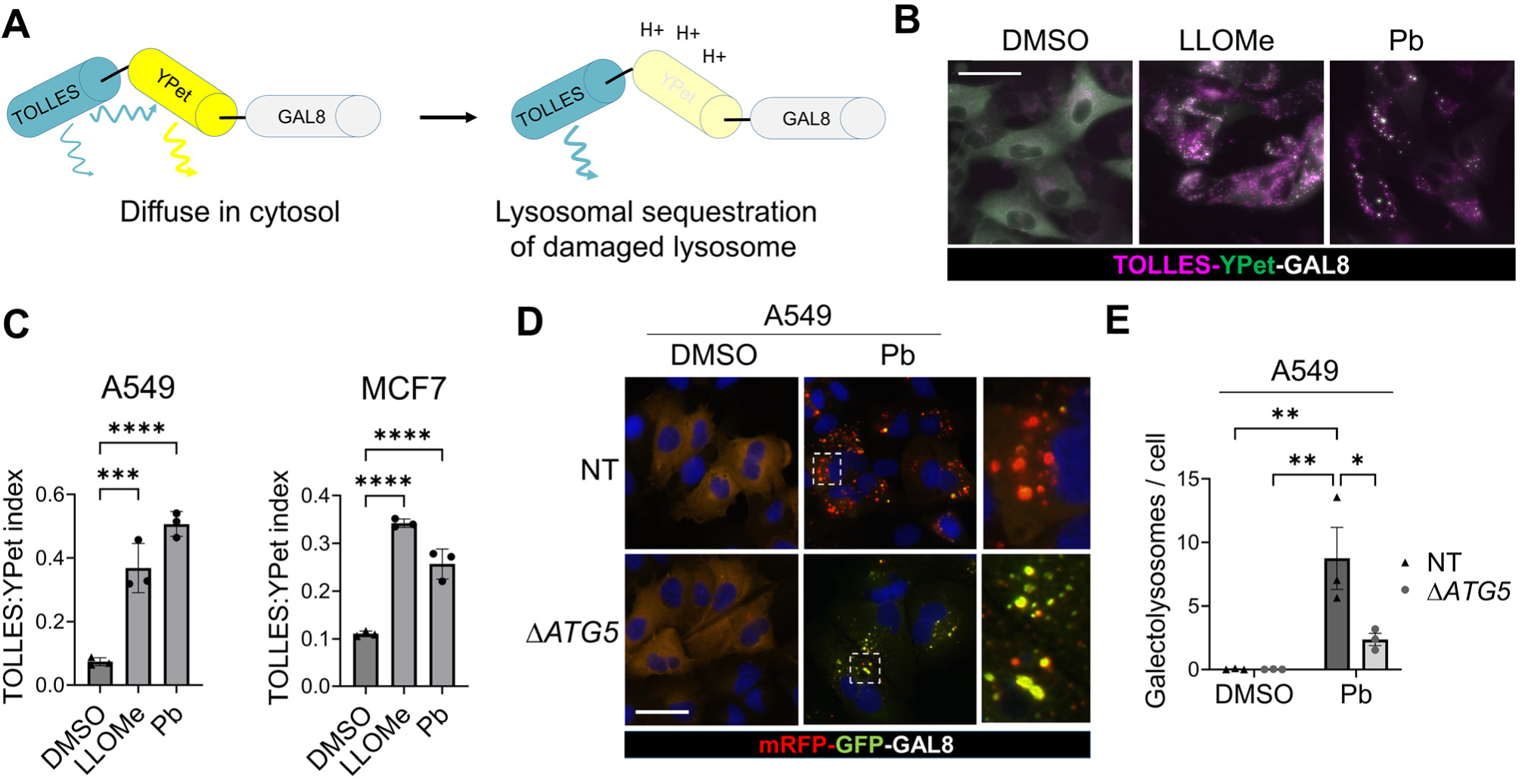
Palbociclib induces lysophagy. **A)** Schematic of SRAI (signal-retaining autophagy indicator)-Galectin 8 (GAL8) reporter for lysophagy. In the cytosol, Ypet emission is low because TOLLES acts as a FRET donor. Conversely, upon completion of lysophagy, Ypet is quenched by low pH and TOLLES fluorescence is apparent. **B-C)** A549- or MCF7-SRAI-GAL8 cells were treated with DMSO, 0.7 mM LLOMe or 10 μM Palbociclib (Pb) for 24 h then imaged by widefield fluorescence. **B)** Representative A549 images (also see **Fig. S1**). **C)** Quantifications of TOLLES:Ypet (lysophagy flux) indices (*n* = 3 independent replicates, mean values ± s.e.m., 50-100 cells per condition per replicate, *** = *P* < 0.001, **** = *P* < 0.0001, 1-way ANOVA , Holm-Šídák comparisons vs. DMSO). **D-E)** A549 NT (non-targeted) or Δ*ATG5* cells expressing mRFP-GFP-GAL8 were treated with DMSO or 10 μM Pb for 24 h prior to imaging by widefield fluorescence. **D)** Representative images, dotted lines demarcate zoom areas. **E)** Quantifications of GFP^-^ mRFP^+^ galectolysosomes (*n* = 3 independent replicates, mean values ± s.e.m., > 90 cells per condition per replicate, ** = *P* < 0.01, * = *P* < 0.05, 2-way ANOVA, Holm-Šídák comparisons). Scale bars = 20 µm.

Next, we examined the steady-state abundance of damaged lysosomes over both acute and chronic treatment regimens. YFP-GAL8 foci, indicative of undegraded and unrepaired damaged lysosomes, were observed in response to chronic 48 h Pb treatment in A549 cells (Fig. 2A), in particular within the autophagy-deficient Δ*ATG5* cells. These YFP-GAL8 foci co-stained with the lysosomal maker lysosomal-associated membrane protein 2 (LAMP2), as expected (Fig. 2B). Quantifying these foci at multiple time points of treatment, it was observed that treatment with Pb provoked a biphasic response in wild-type A549 (NT) cells; a single dose invoked YFP-GAL8 foci accumulation within 1 h, peaking at 24 h, but then resolving by 48 h (Fig. 2C), whereas in Δ*ATG5* cells damaged lysosomes persisted at 48 h. A similar trend was observed for single dose LLOMe treatment, although the peak abundance of damaged lysosomes was observed earlier and, while still evident at 8 h, was completely resolved by 24 h in wild-type (NT) cells (Fig. 2D). Broadly similar responses to Pb and LLOMe were seen in wild-type MCF7 cells, although Palbociclib-induced lysosome damage was still evident at 48 h here, suggesting a lower capacity than A549 cells for *ATG5*-dependent damage resolution (Fig. 2E, F). Finally, the lysosome damaging effect of Pb was observed to be independent of its nucleocytoplasmic targets, CDK4 and CDK6; A549 Δ*ATG5* cells were stably knocked out for CDK4 (Δ*CDK4*) and then CDK6 was acutely knocked out (g*CDK6*), immediately prior to Pb treatment (in order to avoid confounding senescence engaged by double knockout) (Fig. 2G), yet this did not alter YFP-GAL8 focus induction by Pb (Fig. 2H).

**Figure 2.**
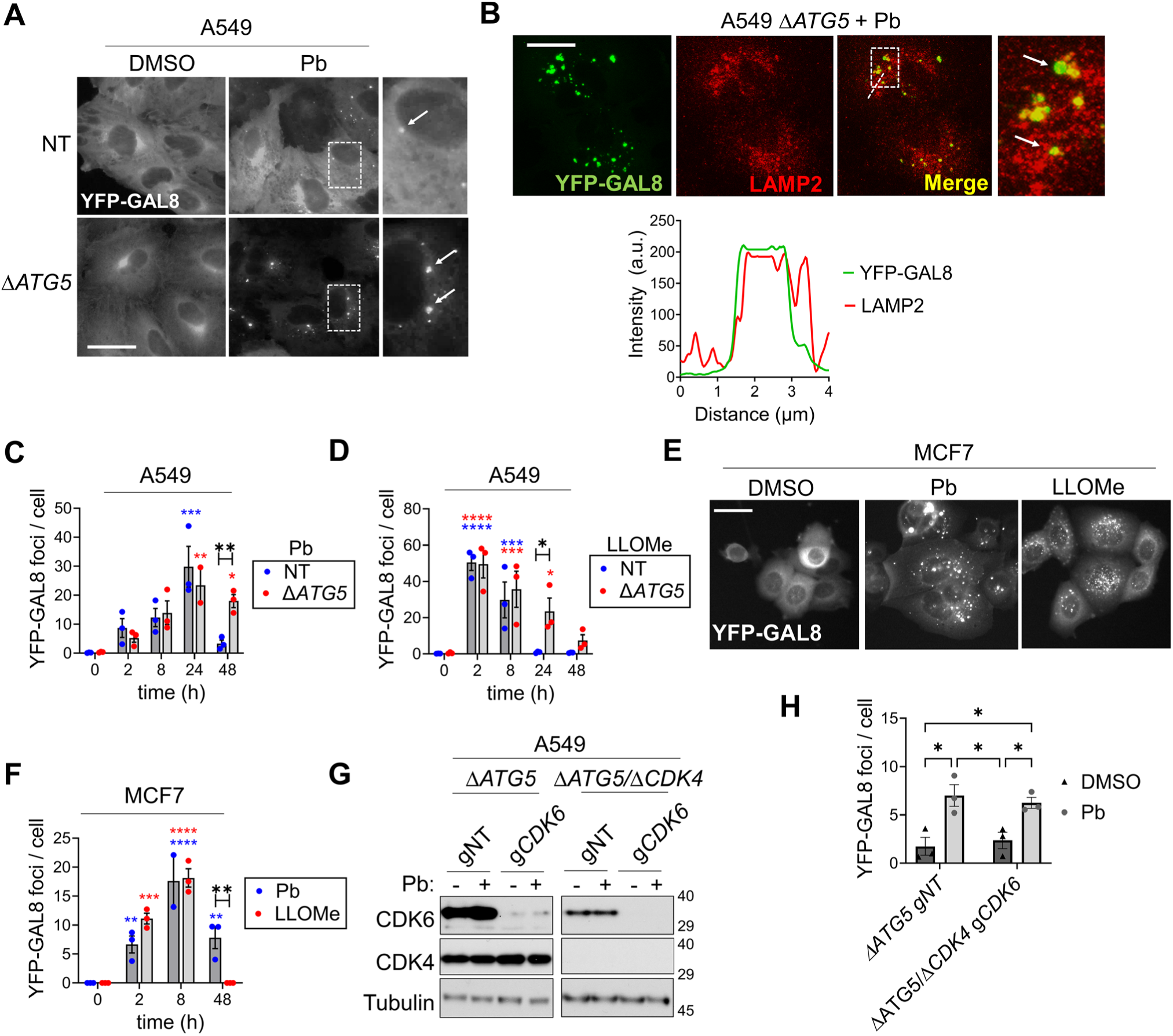
Palbociclib induces chronic lysosome damage. **A-B)** A549 NT or Δ*ATG5* cells expressing YFP-GAL8 were treated with DMSO or 10 μM Pb for 48h. **A)** Representative widefield images, dotted lines demarcate zoom areas, arrows indicate foci. **B)** Representative confocal fluorescence images of LAMP2 co-staining, arrows indicate co-localisation foci in zoom area, dotted line delineates the intensity trace (below). **C-F)** A549 NT, A549 Δ*ATG5* cells and MCF7 cells expressing YFP-GAL8 were treated for indicated time points with 10 μM Pb or 0.7 mM LLOMe, prior to imaging by widefield fluorescence. YFP-GAL8 foci quantifications are presented in **C), D)** and **F)** (*n* = 3 independent replicates, mean values ± s.e.m., > 100 cells per condition per replicate, **** = *P* < 0.0001, *** = *P* < 0.001, ** = *P* < 0.01, * = *P* < 0.05, coloured asterisks indicate significance vs. 0 h for a given treatment, black asterisks indicate significance between different treatments, mixed-effects analysis, Holm-Šídák comparisons). Representative images of MCF7 cells at 24 h shown in **E)**. **G-H)** A549 Δ*ATG5* cells or Δ*ATG5* Δ*CDK4* derivatives, stably expressing Cas9 and YFP-GAL8, were transduced with non-targeting sgRNA (gNT) or g*CDK6*. 72 h post gRNA transduction, cells were treated with DMSO (-) or 10 μM Pb (+), for 24 h and either **G)** immunoblotted or **H)** fixed for widefield fluorescence microscopy quantification of YFP-GAL8 foci, comparing *CDK4/6* intact controls (Δ*ATG5* gNT) with *CDK4/6* double knockout cultures (Δ*ATG5*/gΔ*CDK4* g*CDK6*) (*n* = 3 independent replicates, mean values ± s.e.m., > 110 cells per condition per replicate, ** = *P* < 0.01, * = *P* < 0.05, 2-way ANOVA, Holm-Šídák comparisons). Scale bars = 20 µm.

Considering the above data together, we propose a new role for Pb in inducing chronic lysosome damage and lysophagy, independently of its primary target, CDK4/6. Moreover, the stabilisation of damaged lysosomes in A549 Δ*ATG5* cells, after single treatment with Pb, provides a facile experimental setting for unpicking the events downstream of chronic lysosome damage.

### Damaged lysosomes associate with mitochondria

Under basal conditions, unperturbed lysosomes form transient interorganellar contacts with mitochondria ^51^. Given reports that Pb may also perturb mitochondrial physiology ^52^, we hypothesised that damaged lysosomes could associate with mitochondria. To explore this, we first performed transmission electron microscopy (TEM) of Pb-treated A549 cells (Fig. 3A). Here, particularly in *ATG5*-deficient cells, the presence of swollen, electron-lucent lysosomes was apparent when compared to untreated controls. The increased mass of the lysosomal compartment was corroborated by flow cytometry with Lysotracker Red (LTR) (Fig. 3B) and morphometric quantification of the TEM images (Fig. 3C). The latter analysis also revealed that interorganellar contacts between lysosomes and the mitochondrial network were increased (Fig. 3D, see also arrowheads in zoom panel in Fig. 3A). Thus, we inferred that swollen lysosomes induced by Pb were in contact with the mitochondrial network, although it was not possible to determine if any of these were damaged *per se*. Thus, next, using super-resolution immunofluorescence microscopy, we determined that YFP-GAL8 lysosome foci induced by Pb were indeed associated with — and enwrapped — the mitochondrial network (demarcated by Translocase of Outer Mitochondrial Membrane, TOM20, Fig. 3E, Movie S1). The non-random nature of this association was quantified by analysing the distances of YFP-GAL8 foci from the outer surface of the mitochondrial network and comparing to those observed after YFP-GAL8 foci were computationally shuffled to random positions within the cytoplasmic space (Fig. 3F). Finally, foci dynamics were observed by super-resolution radial fluctuation (SRRF) time-lapse microscopy (Fig. 3G, Movies S2, 3). YFP-GAL8 structures persistently resided along the mitochondrial network, with no or few dissociation events observed over recording times of 10 minutes (Fig. 3G, Movies S2, 3). Taking the above data together, we conclude that chronic damage by Pb results in ruptured lysosomes — or fragments thereof — becoming stably associated with the mitochondrial network.

**Figure 3.**
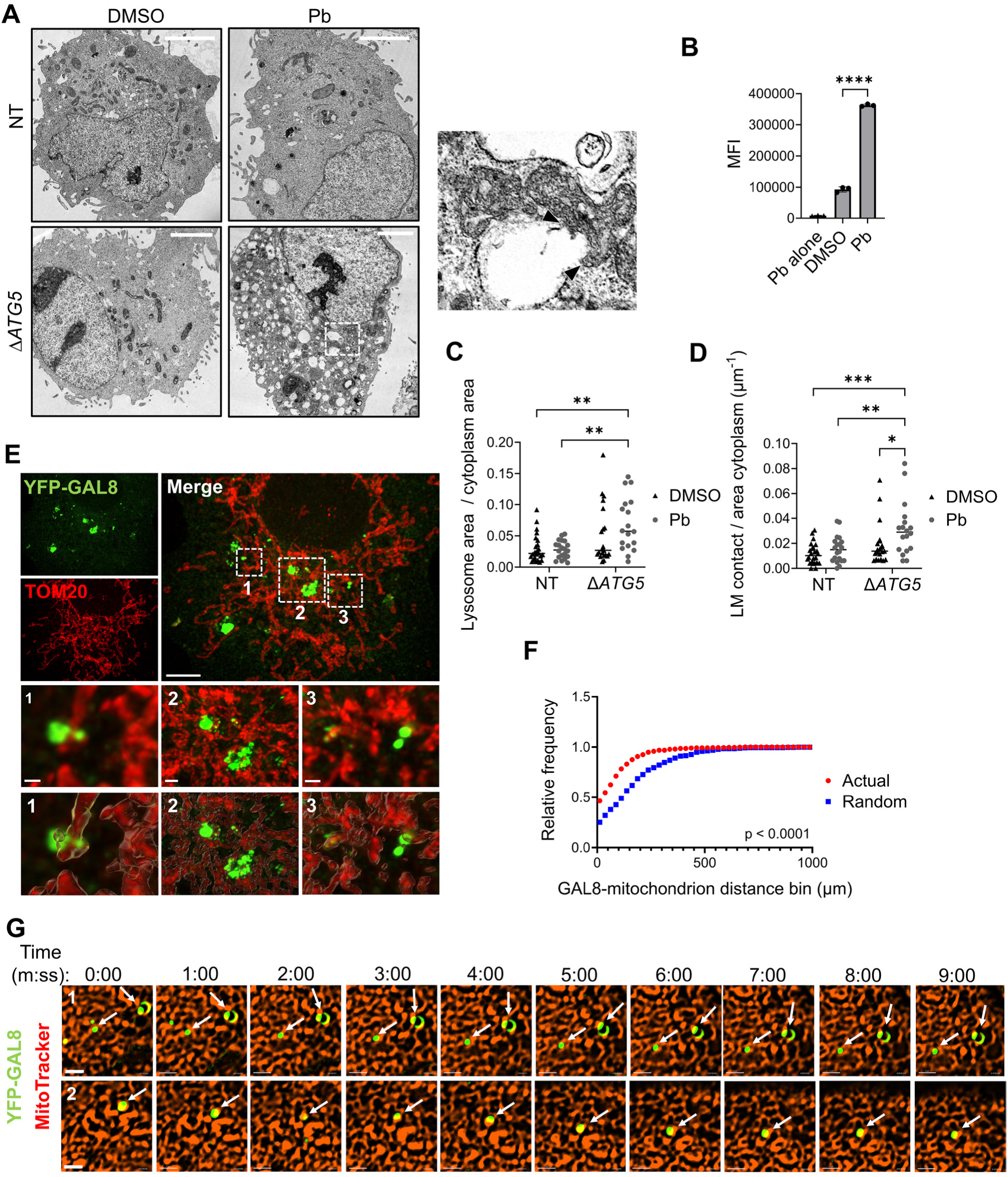
Damaged lysosomes are associated with the mitochondrial network. **A)** A549 NT (non-targeted) or Δ*ATG5* cells were treated with DMSO or 10 μM Palbociclib (Pb) for 48 h then analysed by TEM, representative images shown here (*n* = 20 images per condition). Dotted lines delineate a zoom area showing swollen lysosomes contacting mitochondria (arrowheads delimit a contact site, defined as continuous membrane opposition of < 30 nm). **B)** Δ*ATG5* A549 treated with DMSO or 10 μM Pb for 24 h were incubated for 30 min with Lysotracker Red (LTR, 50 nM) (no LTR in case of Pb alone control). Lysosomal content was analysed by flow cytometry and median fluorescence intensity (MFI) was quantified. (n = 3 independent replicates, mean ± s.e.m., **** = *P* < 0.0001, two-tailed t-test vs DMSO). **C-D)** Quantifications of TEM shown in **A)**. **C)** The proportional cross-sectional area of lysosomes within the cytoplasm. **D)** The length of lysosome-mitochondria (LM) contact sites per unit area of cytoplasm (*n* = 20 images, * = *P* < 0.05, **= *P* < 0.01, *** = *P* < 0.001, 2-way ANOVA, Holm-Šídák comparisons). Scale bars, 0.5 μm. **E-F)** A549 Δ*ATG5* cells expressing YFP-GAL8 were treated with 10 mM Pb for 24 h, then stained for TOM20 (mitochondrion) and imaged by Super Resolution Optical Photon Reassignment Microscopy (SoRA). **E)** Representative raw images in upper panels, dotted lines (1-3) delineate the zoom areas in the row below and the cognate mitochondrial surface rendered images in the bottom row. **F)** Relative frequency distribution of distances between YFP-GAL8 foci and rendered mitochondria (Actual), compared with *in silico* shuffling of these foci within the cytoplasm (Random) (*n* = 400 measurements from 5 cells, Kolmogorov-Smirnov test, *P* < 0.0001, representative of 3 independent experiments). See also **Movie S1**. Scale bars, 5 μm (merge), 1 μm (zooms). **G)** A549 Δ*ATG5* cells expressing YFP-GAL8 were treated with 10 mM Pb for 6 h, then MitoTracker-Red added 30 mins prior to super-resolution radial fluctuation (SRRF) processing-facilitated time-lapse microscopy. White arrows: YFP-GAL8 positive membranes associated with the mitochondrial network. See also: **Movies S2, 3**. Scale bars, 2 mm. Representative of *n =* 4 movies (technical replicates).

### Lysosome damage triggers stereotypical markers of mitochondrial nucleic acid stress

Recently, nucleic acid efflux from the mitochondrion has been described as a signal that triggers interferon responses. This phenomenon frequently correlates with evidence of mitochondrial nucleic acid stress, as visualised by the aggregation of mitochondrial genome nucleoids ^53–55^. We hypothesised that contact between damaged lysosomes and mitochondria might be associated with such mitochondrial nucleic acid stress. Indeed, we observed the formation of large TFAM (mitochondrial transcription factor A)- and DNA-positive foci in the mitochondrial network in response to chronic Pb treatment, indicative of aggregated mitochondrial nucleoids (Fig. 4A, B, C, D). This was greatest in Δ*ATG5* cells, underscoring the relationship between sustained presence of damaged lysosomes and the mitochondrial response to Pb. Corroborating the nucleoid imaging data, similar results were obtained when aggregated nucleoids were alternately imaged by staining for FAST kinase domains 2 (FASTKD2), which marks nucleoid-associated RNA processing granules ^56^ (Fig. 4E, F). Further consolidating the lysosomal association with the mitochondrial stress response, chronic treatment with LLOMe also engendered nucleoid aggregation (Fig. 4G, H). Finally, robust mitochondrial nucleic acid stress responses were also detected in wild-type MCF7 cells treated with Pb or LLOMe (Fig. 4I, J). Taken together, the above data demonstrate that chronic lysosome damage is associated with mitochondrial stress responses.

**Figure 4.**
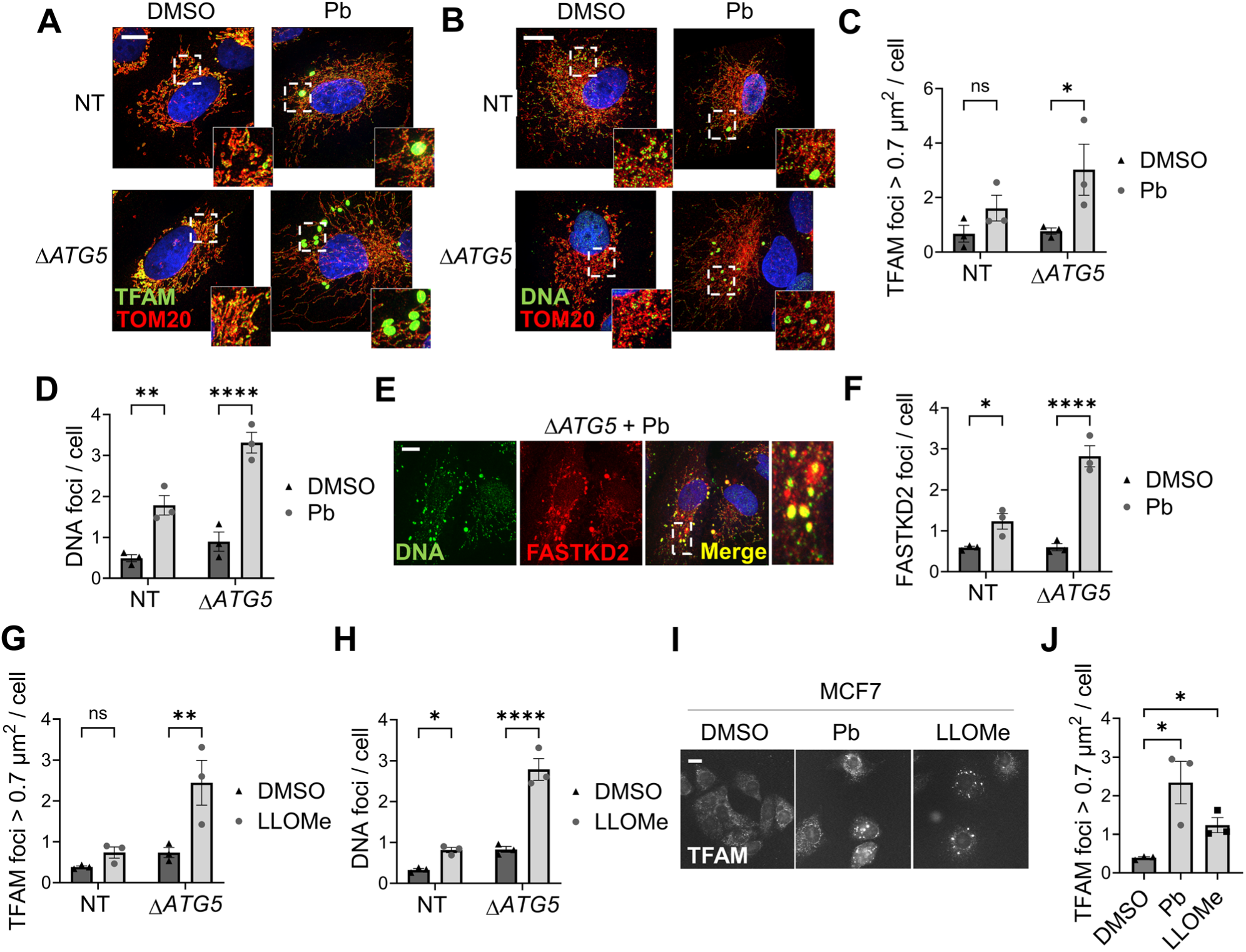
Lysosome damage is associated with stereotypical mitochondrial nucleic acid stress markers. **A-F)** A549 NT (non-targeting wild-type) or Δ*ATG5* cells were treated with DMSO or 10 μM Palbociclib (Pb) for 48 h. Cells were stained for mitochondria (TOM20) and mitochondrial nucleoids using **A,C)** anti-TFAM; **B,D)** anti-DNA; **E,F)** anti-FASTKD2 antibodies. **A,B)** Representative SoRA and **E)** confocal microscopy images. **C-D,F)** Wide-field fluorescence microscopy was used to quantify the aggregated nucleoids (TFAM foci with > 0.7 μm^2^ cross-sectional area or prominent DNA/FASTKD2 foci) (*n* =3 independent replicates, mean values ± s.e.m., > 250 cells per condition per replicate, ns = non-significant, * = *P* < 0.05, **= *P* < 0.01, **** = *P* < 0.0001, 2-way ANOVA, Holm-Šídák comparisons DMSO vs. Pb). Dotted lines delineate zoom areas. **G-H)** A549 NT or Δ*ATG5* cells were treated with DMSO or 700 μM LLOMe for 24 h. Cells were analysed for aggregated mitochondrial nucleoids as in **A)-F)**, quantifications presented here (*n* = 3 independent replicates, mean values ± s.e.m, > 220 cells per condition per replicate, ns = non-significant, * = *P* < 0.05, ** = *P* < 0.01, **** = p < 0.0001, 2-way ANOVA, Holm-Šídák comparisons DMSO vs. LLOMe). **I-J)** MCF7 cells were treated with DMSO, 700 μM LLOMe or 10 μM Pb, for 24 h, then analysed for aggregated TFAM-positive nucleoids as in **A)** and **C). I)** Representative widefield fluorescence images. **J)** Quantifications (*n* = 3 independent replicates, mean values ± s.e.m, > 150 cells per condition per replicate, * = *P* < 0.05, two-tailed t-tests vs. DMSO). Scale bars, 10 μm.

### Palbociclib treatment triggers mitochondrial nucleic acid release into the cytosol and interferon responses

Given the detection of mitochondrial nucleic acid stress upon Pb-mediated lysosome damage, we next sought to directly detect immunostimulatory nucleic acid release from mitochondria. We prepared mitochondrion-free cytosolic extracts (Fig. 5A) from cell clones deleted for Δ*ATG5* or rescued by reintroduction of *GFP-ATG5* ^57^; use of these paired lines diminished potential clonal variation that might have obscured subtle mitochondrial nucleic acid fluctuations. Cytosolic mitochondrial DNA and RNA were found to be elevated in the cytosol in response to chronic Pb; again, this was more prominent in Δ*ATG5* cells where lysosome damage is more poorly resolved (Fig. 5B, C). Given that Pb is known to induce interferon responses, we set out to interrogate whether lysosome damage played a role (and potentially thus cytoplasmic accumulation of mitochondrial nucleic acids). Firstly, we first established that chronic Pb triggers interferon signalling, analysing hallmarks such as phosphorylation of STAT1 (signal transducer and activation of transcription 1) and transcriptional activation of type III interferons, i.e., *IFNL1*, and interferon response genes (IRGs), i.e., *ISG15*, *IFIT1* and *CXCL10*. Pb indeed induced phosphorylation of STAT1 (Figs. 6A, B, S2) and transcriptional activation (Fig. 6C) in A549 cells over 48 h, most prominently in Δ*ATG5* cells, correlating with the persistent lysosome damage known to occur here. Furthermore, phosphorylation of STAT1 was seen when testing lower doses of Pb akin to steady-state plasma concentrations achieved during clinical treatment, although with greater latency (one week) (Fig. 6D). Also corroborating the link with lysosome damage, chronic LLOMe treatment also induced interferon responses over 48 h (Fig. 6E, F). Finally, wild-type MCF7 cells also exhibited interferon responses to chronic treatment with either Pb or LLOMe (Fig. 6G, H, I, J). Taken together, the above data indicate that chronic lysosome damage is a contributor to induction of interferon responses.

**Figure 5.**
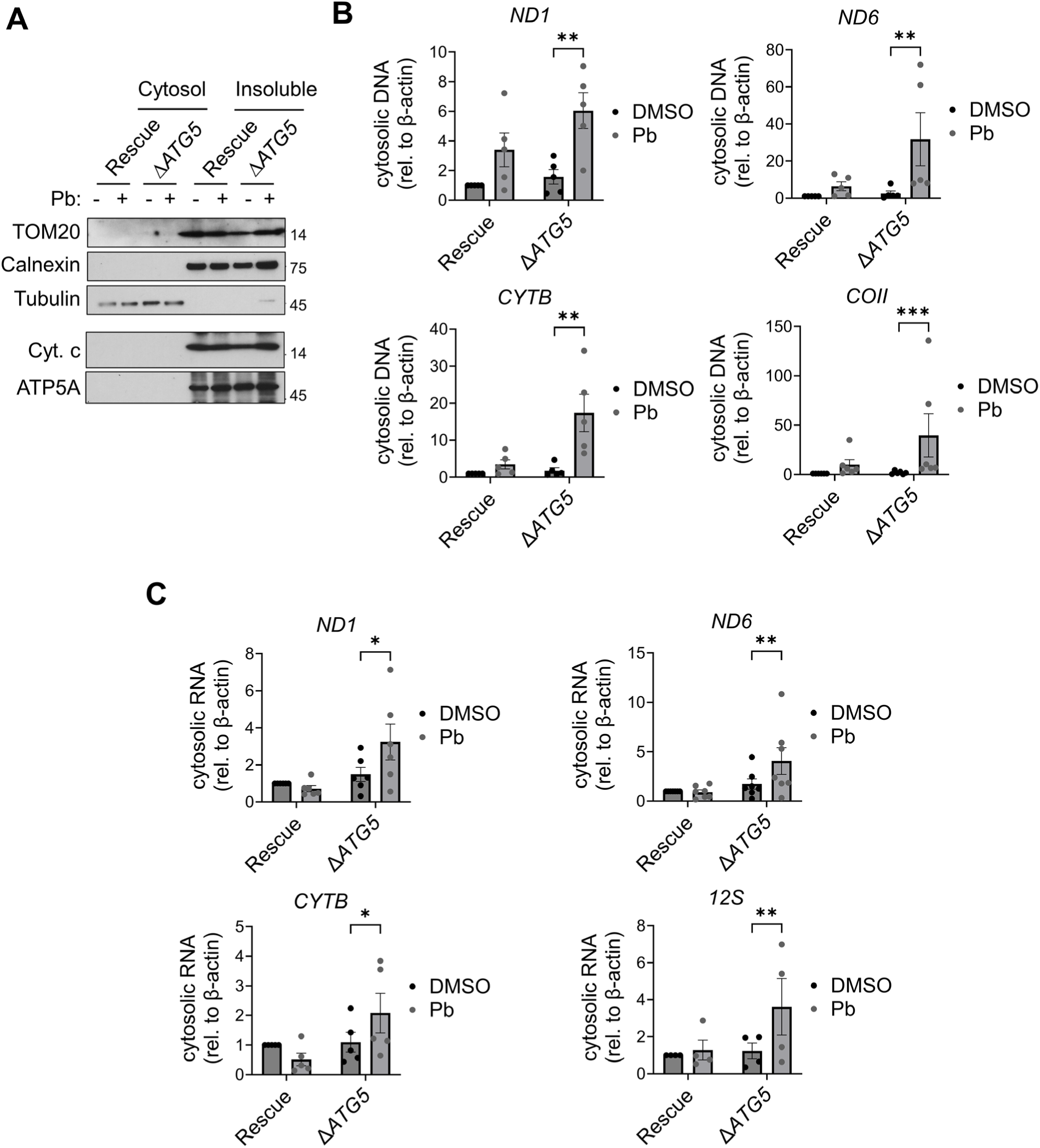
Mitochondrial nucleic acids are released into the cytosol by Palbociclib. **A)** A549 Rescue (Δ*ATG5* + GFP-*ATG5*) and Δ*ATG5* cells were treated with DMSO (-) or 10 μM Palbociclib, Pb (+), for 48 h. Pure cytosolic fraction (Cytosol) was separated from remaining cellular material including mitochondria (Insoluble) using digitonin extraction and immunoblotted to confirm purity (mitochondrial markers: TOM20; Cytochrome C; ATP5A, ER marker: Calnexin). Cytochrome C and ATP5A are from a second immunoblot on same samples from the same replicate. Representative of *n* = 3 independent replicates. **B-C)** Nucleic acids (**B**, DNA; **C**, RNA) were isolated from the cytosolic fraction and qPCR performed for mitochondrial genes *ND1*, *ND6*, *CYTB* and *COII*, normalised to *ACTB* from whole cell DNA (*n* = 4 independent replicates, minimally, after outliers removed by Grubb’s test, mean ± s.e.m., normalisation to Rescue + DMSO, ** = *P* < 0.01, *** = *P* < 0.001, two-tailed ratio paired t-tests).

**Figure 6.**
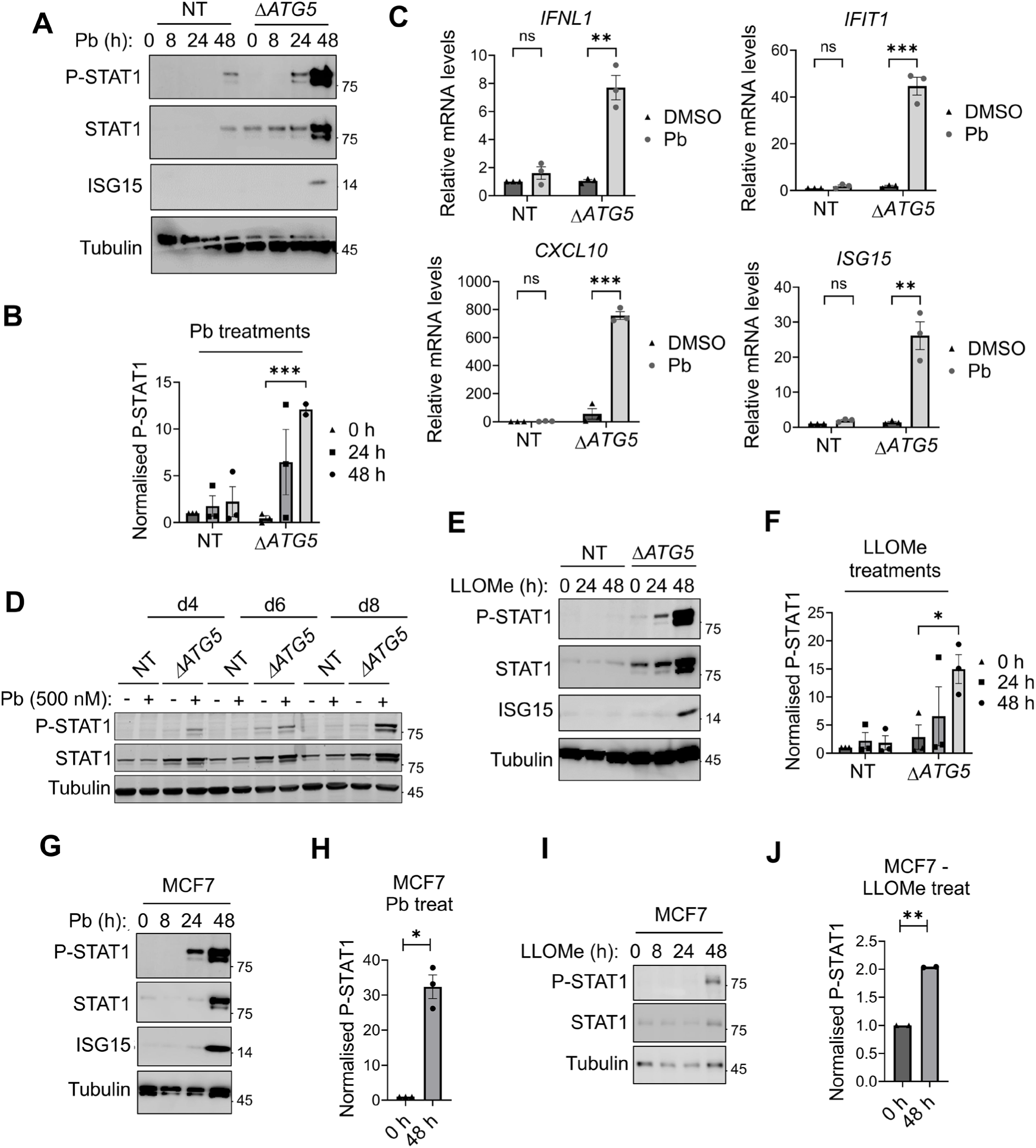
Chronic lysosome damage triggers interferon responses. **A-B)** A549 NT (non-targeted wild-type) or Δ*ATG5* cells were treated with 10 μM Palbociclib (Pb) for the indicated time and then immunoblotted for interferon pathway activation (phospho-Y701 STAT1, P-STAT1, and ISG15). **A)** Representative blot. **B)** Quantifications (*n* = 3 independent replicates, mean ± s.e.m., outlier removed by Grubb’s test, *** = *P* < 0.001, 2-tail t-tests vs. 0 h, Holm-Šídák correction). **C)** A549 NT or Δ*ATG5* cells treated as in **A** and subjected to qRT-PCR for indicated genes (*n* = 3 independent replicates, mean ± s.e.m., relative to *18S*, normalised to NT + DMSO, ns = non-significant, ** = *P* < 0.01, *** = *P* < 0.001, 1-sample t-test for NT, 2-tailed t-test for Δ*ATG5*). **D)** A549 Δ*ATG5* cells were treated with DMSO (-) or 500 nM Pb (+), for the indicated number of days (d), then immunoblotted for P-STAT1 (*n* = 3 independent replicates, representative replicate shown). **E-F)** A549 NT (non-targeted wild-type) or Δ*ATG5* cells were treated with 700 μM LLOMe for the indicated times then immunoblotted for interferon pathway activation (P-STAT1, ISG15). **E)** Representative blot. **F)** Quantifications (*n* = 3 independent replicates, mean ± s.e.m., * = *P* < 0.05, 2-tail t-tests vs. 0 h, Holm-Šídák correction). **G-J)** MCF7 cells were treated with 10 μM Palbociclib (Pb) (**G**-**H**) or 700 nM LLOMe (**I**-**J**) for the indicated times then immunoblotted as shown. **G,I)** Representative blots. **H,J)** Quantifications (*n* = 3 independent replicates, mean ± s.e.m., * = *P* < 0.05, ** = *P* < 0.01, 1-sample t-tests).

### Mitochondrial nucleic acids are required for the interferon response to chronic lysosome damage

To test whether the observed release of mitochondrial nucleic acid triggers the interferon signalling associated with Pb-induced lysosome damage, we next depleted mitochondrial genomes from A549 control (Rescue) and Δ*ATG5* cells, via chronic dideoxy-cytidine treatment, generating “Rho0” cultures (Fig. S3A). Importantly, mitochondrial nucleic acid depletion inhibited engagement of downstream markers of interferon production by Pb, including phosphorylation of STAT1 (Fig. 7A, B), and transcriptional upregulation of *IFNL1* and *ISG15* (Fig. 7C). Two modes of nucleic acid efflux from mitochondria have been postulated in the prior literature. The first is associated with stereotypical markers of mitochondrial nucleic acid stress, as observed in Fig. 3, and can be blocked using the VDAC channel inhibitor VBIT-4 ^53–55^. The second is characterised by a requirement for mitochondrial BAK (BCL2 antagonist/killer 1) and BAX (BCL2-associated X, apoptosis regulator) pore-forming proteins ^58,59^. We found that interferon responses to Pb were indeed abrogated by VBIT-4 treatment (Fig. 7D, E) but not *BAK* and *BAX* double knockout (Fig. S3B). This observation further corroborates the link between the mitochondrial nucleic acid stress induced by lysosome damage and interferon signalling. In line with this, chronic LLOMe treatment was also unable provoke phosphorylation of STAT1 in Rho0 cells (Fig. 7F, G). Finally, given the association of mitochondrial depolarisation with at least some forms of mitochondrial perturbation, we assessed this; however, Δ*ATG5* cells did not show depolarisation as assessed by flow cytometry with JC-10 dye, whereas they did respond in this manner to a positive control drug, GTTP (Gamitrinib-triphenylphosphonium) ^60^ (Fig. 7H). Taken together with our prior findings, the above data demonstrate that lysosome damage triggers accumulation of mitochondrial nucleic acid in the cytosol, and that this drives interferon responses. However, Pb also reportedly triggers interferon responses by suppressing CDK4/6-mediated silencing of nuclear genome-derived dsRNA ^34^. Given that we observe that CDK4/6-independent lysosome damage is associated with mitochondrial nucleic acid release, we hypothesised co-operation between on-target (kinase activity) and off-target (lysosomal-mitochondrial) effects of Pb. Indeed, *CDK4/6* deletion, while insufficient to engage robust interferon responses alone, synergised with Pb treatment to induce a maximal response (Fig. 7I, J). Taking the above data together, we conclude that sustained suppression of CDK4/6 kinase activity has the potential to engage interferon signalling via non-mitochondrial routes ^34^, but that the lysosome-mitochondrion axis must co-operate to fully realise the interferon effect, at least in the context of Pb treatment.

**Figure 7.**
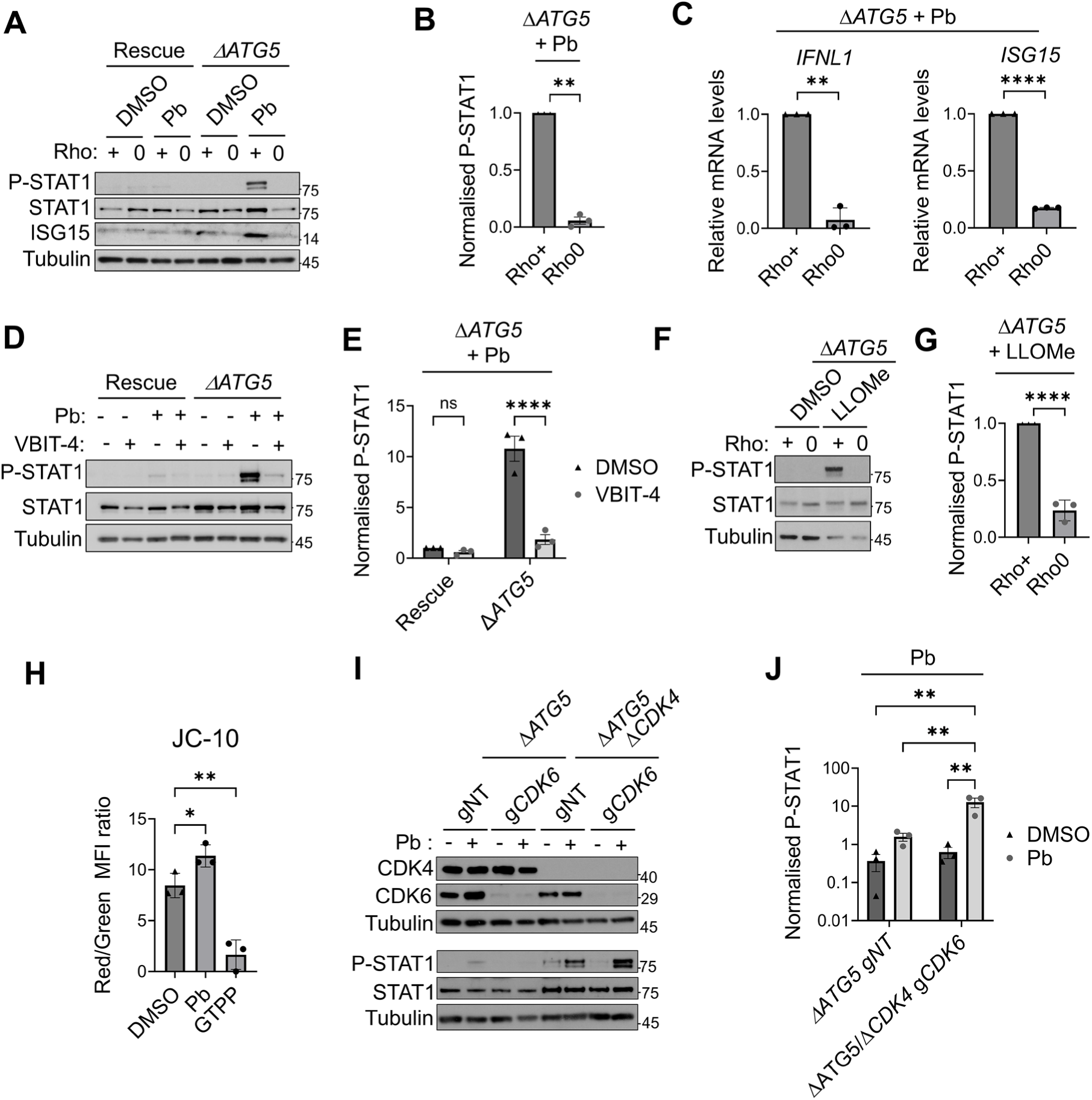
Mitochondrial nucleic acids are required for interferon responses associated with chronic lysosome damage. **A-C)** A549 Rescue (Δ*ATG5* + GFP-*ATG5*) and Δ*ATG5* cells were treated to generate control (Rho+, +) or mitochondrial genome depleted (Rho0, 0) cells. Cells were treated with DMSO or 10 μM Palbociclib (Pb) for 48 h and immunoblotted for interferon activation (phospho-Y701-STAT1, P-STAT1, ISG15). **A)** Representative blot. **B)** Quantification for Pb-treated Δ*ATG5* cells (*n* = 3, mean ± s.e.m., normalised to Rho+, ** = *P* < 0.01, ns = non-significant, 1-sample t-test). **C)** A549 Δ*ATG5* cells (Rho+ or Rho0) were treated with 10 μM Pb for 48 h and qRT-PCR performed for *IFNL1* and *ISG15* (*n* = 3 independent replicates, mean ± s.e.m., relative to *18S*, normalised to Rho+, ** = *P* < 0.01, **** = *P* < 0.0001, 1-sample t-tests). **D-E)** A549 Rescue or Δ*ATG5* cells were treated with DMSO or 10 mM Pb +/- 15 mM VBIT-4, then immunoblotted as shown. **D)** Representative blot. **E)** Quantification for Pb-treated Δ*ATG5* cells (*n* = 3 independent replicates, mean ± s.e.m, ns = non-significant, **** = *P* < 0.0001, 2-way ANOVA, Holm-Šídák comparisons DMSO vs. VBIT-4). **F-G)** A549 Δ*ATG5* cells, Rho+ (+) or Rho0 (0), were treated with DMSO or 700 μM LLOMe for 48 h, then immunoblotted for P-STAT1. **F)** Representative blot. **G)** Quantification for LLOMe treated cells, normalised to Rho+ (*n* = 3 independent replicates, mean ± s.e.m., **** = *P* < 0.0001, 1-sample t-test). **H)** Δ*ATG5* A549 were treated with DMSO, Pb (10 μM, 24 h) or Gamitrinib TPP (GTPP, 20 μM, 4 h). Cells were then incubated for 30 min with JC-10 (15 μM) prior to analysis by flow cytometry. Quantitative fluorescent ratio of JC-10 (aggregating (red, em 585) versus monomeric (green, em 525)) was used to assess mitochondrial membrane potential (MFI, median fluorescence intensity) (*n* = 3 independent replicates, mean ± s.e.m., ** = *P* < 0.01, * = *P* < 0.05, 1-way ANOVA, Holm-Šídák comparisons vs. DMSO). **I-J)** A549 Δ*ATG5* cells or Δ*ATG5 ΔCDK4* derivatives, stably expressing Cas9, were transduced with non-targeting sgRNA (gNT) or g*CDK6*. 72 h post transduction, cells were treated with DMSO (-) or 10 μM Pb (+) for 24h, then immunoblotted as shown. **I)** Representative blot. **J)** Quantifications comparing CDK4/6 intact controls (Δ*ATG5* gNT) with CDK4/6 double knockout cultures (Δ*ATG5/ΔCDK4* g*CDK6*) (*n* = 3 independent replicates, mean ± s.e.m., ** = *P* < 0.01, 2-way ANOVA, Holm-Šídák comparisons).

## Discussion

Our findings support the following model (Fig. 8). In response to chronic exposure to the lysosomotropic cancer therapeutic Palbociclib (Pb), persistent lysosome rupture is invoked. Acute lysosome damage typically leads to repair, removal and/or replacement of the damaged organelle. These responses are at the core of the ELDR. We propose that in cells where chronic lysosome damage is insufficient to trigger cell death, propagation of stress signalling via lysosome-mitochondrion association leads to accumulation of cytosolic, self-derived immunostimulatory nucleic acids, triggering interferon signalling. Indeed, other recent data have hinted that stress signals emanating from acutely damaged lysosomes trigger wider changes in cell physiology ^30–32^. However, we acknowledge that – despite the triggering of such mitochondrial phenomena also by the tool compound LLOMe – it is possible that direct effects of Palbociclib on mitochondria contribute to this phenotype.. It is also possible the involvement of other organelles that form contacts with both mitochondria and lysosomes – such as the endoplasmic reticulum (ER) ^61^ – could be involved in lysosome-mitochondrial communication, but this remains to be tested.

**Figure 8.**
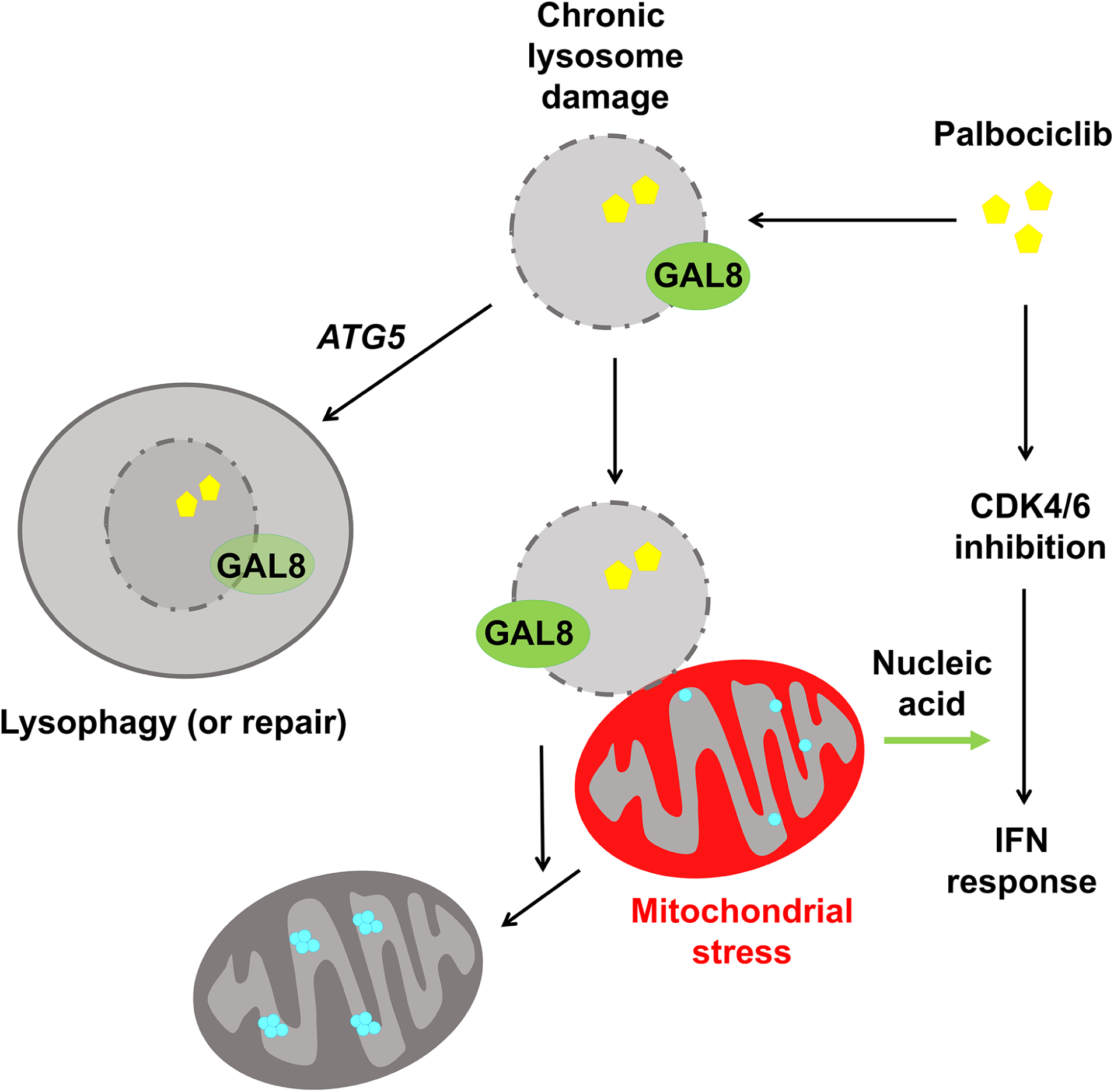
Model for response to chronic lysosome damage invoked by Palbociclib. See text for details. GAL8 = galectin 8. IFN = interferon. Blue dots depict mitochondrial nucleoids.

In the case of Pb treatment, the mitochondrial response associated with lysosome damage is required for interferon signalling but maximal stimulation occurs in co-operation with the known CDK4/6-inhibitory activity of the drug, explaining fully this important anti-cancer facet of Pb action. Instructively, mitochondrial nucleic acids may similarly be required to license interferon responses after *Salmonella* infection ^62^. This is suggestive when one considers that bacterial phagosome damage by infectious agents can also engage selective autophagy mechanisms with molecular parallels to lysophagy ^2^. Furthermore, it has been recently suggested that damage to lysosomes invoked by pathogens effects broad changes in mitochondrial proteome, secondarily to vesicular rupture ^63^. Taking these observations together, we speculate that mitochondrial responses associated with lysosome damage may have initially evolved to consolidate infection responses by priming interferon responses, but not in itself triggering maximal signalling, i.e., co-operating with additional stimuli such as nucleic acids or other patterns physically derived from the infectious agent. However, these ideas require future exploration.

Our data show that the lysosome damage-induced interferon response depends on chronic perturbation. For instance, A549 cells deleted for Δ*ATG5*, which are deficient in the ELDR, show a more robust response to Pb and LLOMe, which correlates with more sustained lysosome damage. Similarly, wild-type MCF7 cells show evidence of resolution of lysosome damage but more slowly than wild-type A549 cells, and this again correlates with a more readily detectable interferon response. However, it is possible that other functions of *ATG5* in autophagy pathways, for instance degradation of PRRs, could also contribute to the observed *ATG5-*dependency in A549 cells ^64–67^. Similarly, the loss of mitophagy in these *ATG5*-deficient cells could also contribute to high levels of interferon response ^68^. Overall, our data nonetheless still establish an association between lysosome damage and engagement of interferon responses via mitochondrial release of nucleic acid.

Finally, our results highlight an important consideration when investigating small-molecule therapeutics for cancer. The lysosome is sometimes considered a passive problem for therapy, in that it sequesters basophilic drugs from their site of action ^33^. Although the mitochondrion is implicated here in response to Pb, it is unlikely that it will ultimately prove to be the only consequential signalling hub responding to chronic lysosome damage. Indeed, we would suggest that off-target signalling effects of cancer therapeutic drugs consequent from chronic accumulation in the lysosome should be factored into cell-based drug action studies in future. This will facilitate holistic understanding of therapeutic compound action and thus, ultimately, promote refinement of clinical applications.

## Acknowledgements

We thank Richard Youle, Viktor Korolchuk, Noor Gammoh, Felix Randow for reagents. We thank Noor Gammoh and Alex von Kriegsheim for feedback. We thank Steven Mitchell for assistance with electron microscopy. For the purpose of open access, the author has applied a CC-BY public copyright licence to any Author Accepted Manuscript version arising from this submission.

## Funding

This work was supported by a Cancer Research UK (C20685/A29576 to S.W.).

## Methods

### Cell culture and cell lines

Cells were cultured in DMEM supplemented with 10% FBS, 2 mM L-glutamine and 10 U ml^-1^ 10 μg ml^-1^ penicillin/streptomycin, at 37°C under 5% CO_2_. At monthly intervals all cells in culture were tested for the presence of mycoplasma. A549 NT, D*ATG5* and Rescue (D*ATG5* + GFP-ATG5) lines were described previously by the authors and are a derivative of the A549:EcoR parental cell line, which is A549 human lung adenocarcinoma cells derivatised to express Ecotropic receptor to enable transduction with retrovirus with murine tropism ^69^. MCF7:EcoR cells were newly generated, by the same protocol, in this study, from parental MCF7 cells. Parental A549:EcoR and MCF7 breast carcinoma lines were from the authors’ laboratory and were verified by microsatellite genotyping. YFP-GAL8 expressing retrovirus (see **Plasmids**) was used to generate stable A549 line pool, as described previously ^69^; pools of transductants were selected with 10 mg ml^-1^ blasticidin for 10 days. Stable MCF7 cell pools expressing YFP-GAL8 were similarly generated from parental MCF7-EcoR cells (selection in 50 μg ml^-1^ blasticidin). A549-EcoR cells stably expressing mRFP-GFP-GAL8 were made by lentivirus transduction (see **Plasmids**) and a pool of transductants selected with 10 mg ml^-1^ blasticidin. A549-EcoR and MCF7 cells stably expressing SRAI-GAL8 were created by lentiviral transduction (see **Plasmids**) and selection with 10 mg ml^-1^ blasticidin. Then, cell populations expressing high YPet and TOLLES fluorescent levels were enriched using a FACSAria II (BD Bioscience, San Jose, CA). To enable both pooled and single clone CRISPR/Cas9 knockouts in A549-EcoR cell lines, stable expression of Cas9 was achieved by transducing lentivirus made from LentiCas9-Blast template and selection with 10 mg ml^-1^ blasticidin. CRISPR/Cas9-mediated knockout in the resulting lines expressing Cas9 was performed using lentiviral delivery of gRNA. Non-targeting/targeting sgRNA was cloned into lentiGuide-puro for lentivirus production, and transductants selected for with 2.5 μg ml^-1^ puromycin. gRNA sequences were: GFP (NT): GTGAACCGCATCGAGCTGAA; Luciferase (NT-2): AACGGCGGCGGGAAGTTCAC; MAVS: TCTTCAATACCCTTCAGCGG; CDK4: TGTAGTGATGCGCGACGACA; CDK6: CTATTACTGAGCTCGATGTG; BAX: GGCGGTGATGGACGGGTCCG; BAK: GGAACTCTGAGTCATAGCGT. Where single cell clones were used, this was achieved via limiting dilution. All lentivirus was packaged in HEK293T cells according to standard protocols; briefly, viral vector was co-transfected with pMD2.G and psPAX2 packaging plasmids using Lipofectamine 2000.

### Drug treatments

Seeding density was normalised per experiment treatment requirements. Cells were plated to be approaching confluency, but not yet confluent, at the point of analysis. Typically, one third of the number of cells to be treated with higher doses (e.g. 10 μM) Pb (Pb-HCL, Cayman #16273, dissolved in DMSO) were seeded relative to the cognate DMSO vehicle treatments for 48 h treatments, and one half for 24 h treatments, to compensate the cytostatic effect of Pb. For 10 mM Pb treatment, treatment was applied once for the entire duration. For longer term treatments using 500 nM Pb, cells were plated on day 0, treatment was initiated on day 1, and medium was refreshed with new drug or vehicle at day 4. LLOMe (LLOMe-HCL, Cayman #16008, dissolved in DMSO) was dosed at 700 mM. VBIT-4 (Selleckchem #S3544) was dissolved in DMSO and used at 10 mM.

### Mitochondrial genome depleted cultures

To generate paired Rho^+^ and Rho0 cultures, cells were cultured in standard medium supplemented with 200 μM uridine (Sigma #U3750) and, for Rho0 culture, with an additional 10 μM ddC (2’,3’-dideoxycytidine, Sigma #D5782) for 14 consecutive days ^48^. Cells were then maintained in culture for a maximum of another week in medium still supplemented with ddC.

### Plasmids

See Table S1 for full list. Where Gateway cloning was performed this was as per standard protocols using BP clonase II and LR clonase II (Invitrogen). sgRNA cloning into lentiGuide-puro vector was performed using restriction digestion as per the standard protocols available from Addgene. Detailed sequence maps are available from authors upon request.

### Immunoblotting

Cells were lysed in 4% SDS lysis buffer (4% SDS, 50 mM Tris-HCl pH 7.5 and 150 mM NaCl), followed by pulse sonication. Protein concentration was measured by BCA protein assay, using the Pierce™ BCA protein assay kit (Thermo Fisher, #23225). Normalised protein samples were diluted in Laemmli buffer (4% SDS, 20% glycerol, 10% 2-mercaptoethanol, 0.125 M Tris-HCl pH 6.8, bromophenol blue) and briefly heated to 95°C. SDS-PAGE was carried out using either 4-12% NuPage™ Bis-Tris Precast mini, or 4-12% Novex™ Tris-Glycine precast midi gels (Thermo Fisher), in 1x NuPage™ MOPS SDS running buffer (Thermo Fisher) or Pierce™ Tris-Glycine SDS running buffer (Thermo Fisher), respectively. Wet transfer was used to transfer protein from mini gels, performed onto 0.2 mm nitrocellulose membrane (GE Life Sciences), and semi-wet transfer system (Trans-blot turbo, Bio-Rad) was used to transfer proteins from midi gels. Membranes were blocked in 5% BSA (Sigma) in PBS for 1 h at room temperature (RT). All washes were performed three times in TBS - 0.1% Tween 20 (TBST), for 10 min each at RT. Primary antibody dilution was in 0.1% BSA in PBS with 0.05% sodium azide, typically at 1:1000 except for α-Tubulin (1:50000). Membranes were incubated in primary antibody solution overnight at 4 °C. The following antibodies were used: anti-ATG5 (Sigma, #A0731), anti-Calnexin (Cell Signalling Technology [CST], #2679), anti-CDK4 (CST, #12790), anti-CDK6 (CST, #3136), anti-ISG15 (CST, #2743), anti-MAVS (CST, #3993), anti-phosho-Y701-STAT1 (CST, #9167), anti-STAT1 (CST #9176), anti-TOM20 (Santa Cruz, #sc-17764), anti-Cytochrome C apoptosis antibody cocktail (Abcam, #ab110415) and anti-α-Tubulin (Sigma, #T9026). Secondary antibody solution was 0.1% powdered milk in TBST in 1:3000 dilution. Membranes were incubated with cognate HRP-linked secondary antibody (CST) for 1 h at RT. For detection, membranes were incubated in ECL reagent (ECL prime, Western blotting reagent, GE Life Sciences) for 5 min at RT, or ECL reagent (1 M Tris-HCl pH 8.5, 250 mM luminol, 90 mM coumaric acid and 0.002% H_2_O_2_) for 1 min at RT, followed by film exposure and development (AGFA automated developer), or chemiluminescent imaging (Amersham ImageQuant 800, software v1.2.0). Relative protein band intensities were quantified using FIJI (v2.1.0). For P-STAT1, this is given as a ratio to tubulin, as STAT1 itself is an interferon stimulated gene. Uncropped blots are shown in Figs. S4-6.

### Immunostaining

For immunofluorescence experiments, cells were typically seeded onto round 16 mm cover glass (Fisher Scientific) except for super-resolution microscopy where cells were seeded on high-precision cover glass (Zeiss™, Fisher Scientific). Cells were fixed in 4% paraformaldehyde (PFA) / PBS for 10 min at RT and washed once in PBS. Permeabilization was performed for 10 min at RT using 0.25% Triton X-100 (Sigma) / PBS, followed by another PBS wash. Primary and secondary antibody solution was diluted in PBS / 1% BSA, typically at 1:200 (except p62 used at 1:1000). Secondary antibody solution typically contained DAPI at 1:2500. Cells were incubated with primary antibody solution for 1 h, at RT, followed by 3 x 5 min PBS washes. The following antibodies were used: anti-DNA (EMD Milipore, #CBL186), anti-FASTKD2 (Proteintech, #17464-1-AP), anti-LAMP2 (Abcam, #ab25631), anti-TFAM (CST, #8076), anti-TOM20 (CST, #42406) (used when co-staining with mouse antibodies), anti-TOM20 (Santa Cruz, #sc-17764) (used when co-staining with rabbit antibodies and YFP-GAL8) and anti-DNA (Merck-Millipore, #CBL186). Secondary antibody was cognate fluorophore linked secondary antibody (Thermo Fisher, Alexa Fluor). Secondary antibody solution was applied to cells for 45 min, at RT, followed by 3 x 5 min PBS washes. Regular cover glass was mounted onto a microscope slide using DAKO fluorescence mounting medium. High-precision cover glass super-resolution microscopy was mounted using Fluoromount™ aqueous mounting medium (Sigma).

### Fluorescence Microscopy

For widefield immunofluorescence microscopy, images were taken with Olympus BX51 fluorescent microscope (Cell^F software). For confocal microscopy, images were taken using a Nikon A1R laser-scanning confocal microscope. Images are presented as maximum projection Z-stacks. Super-resolution images were acquired using Super Resolution Optical Photon Reassignment Microscopy on a Nikon Eclipse Ti2 inverted microscope stand and a SR Apo TIRF 100x 1.49NA oil lens (Nikon Instruments). The CMOS cameras used for acquisition were Teledyne Photometrics Prime 95B (Teledyne Photometrics 3440 E.Britannia Drive, Tucson AZ) and 405, 488, 561, 640nm laser lines. Acquisition of images and deconvolution was carried out using Nikon NIS-Elements Advanced Research software (v5.21-03). For imaging SRAI-GAL8, cells were plated on glass bottom 96 well plates (P96-1.5H-N, Cellvis) at 70-90% confluency. Cells were fixed with 4% paraformaldehyde in PBS, pH 7.2 at room temperature for 10 mins. Cells were then washed twice with PBS and kept at 4°C. Cells were imaged 1-3 days after fixation while immersed in PBS. Images were captured with a Nikon Eclipse Ti-E inverted microscope (Nikon Instruments UK Ltd, Kingston Upon Thames, UK) using a 40X objective. Widefield illumination for this was provided by a CoolLED PE-4000 LED light source (CoolLED Ltd, Andover, UK) combined with LED-CFP/YFP/mCherry-3X-A (Pinkel) filter sets (IDEX Health & Science, LLC, Center of Excellence, 1180 John Street, Rochester, New York 14586, United States). Images were acquired with a Photometrics Prime BSI sCMOS camera (Teledyne Photometrics, 3440 E. Britannia Drive, Suite 100, Tucson, AZ 85706). Conditional imaging was setup and controlled using the Nikon Nis-Elements JOBS module. The JOB (script) incorporates automated two wavelength capture of 3D images (Z stacks for CFP and YFP filters) from an array of imaging sites (XY positions) using a motorised XY stage within user selected wells (a total of 7 fields per well were imaged). SRAI-GAL8 images are shown as z-projections from 5 z-steps.

### SRRF Time-lapse videomicroscopy

Time-lapse videomicroscopy was performed by exchanging full medium for phenol-red free medium containing 0.132 μg ml^-1^ MitoTracker Red and fresh Pb 30 mins prior to imaging. Images were acquired on the multimodal Imaging Platform Dragonfly (Andor technologies, Belfast UK) equipped with 405, 445, 488, 514, 561, 640 and 680nm lasers built on a Nikon Eclipse Ti-E inverted microscope body with Perfect focus system (Nikon Instruments, Japan). Super Resolution Radial Fluctuations (SRRF) data were acquired using the iXon 888 EMCCD camera and a 100x SR Apo TIRF 1.49NA objective lens. Environmental control of the cultured cells was maintained during imaging with an Okolab bold line stage top incubation chamber incorporating temperature and humidified CO2 control (Okolab S.R.L, Ottaviano, NA, Italy).

### Transmission electron microscopy (TEM)

Fixative solution was prepared by mixing fresh aqueous 16% paraformaldehyde (from methanol-free powder, Sigma) and fresh 25% EM grade glutaraldehyde (Merck) to final concentration 2% paraformaldehyde, 2.5% glutaraldehyde in 0.1 M PHEM buffer (120 mM PIPES, 50 mM HEPES, 4 mM MgCl_2_ and 20 mM EGTA, pH 6.9). Two 15 cm dishes per condition of cells were used and cells pelleted by centrifugation 200 g for 3 min in a 15 mL tube. Cells were resuspended in the fixative solution and incubated for 20 min at room temperature, on a slow-motion roller bank. Cells were then centrifuged, supernatant removed, and fresh fixative solution added. The second fixative solution was incubated with cells for 4 h at room temperature, on a slow-motion roller bank. The cells were washed 3 times with 0.1 M PHEM buffer and stored in 0.1 M PHEM until embedding in EPON resin at 4°C (2-3 days). Sections were co-stained with uranyl acetate and lead citrate to provide membrane contrast. TEM images were taken by JEOL JEM-1400 series, 120 kV transmission electron microscope.

### Image analyses

From SRAI-GAL8 images, a TOLLES:YPet index was obtained using an ImageJ macro that calculates the percent area of pixels with TOLLES:YPet ratio above a given ratio over the total area that the fluorophore occupies within the field ^50^. A ratio of 1.75 was used. Substructures in cells (foci, lysosomes, nuclei) were generally counted manually using ImageJ cell counter plug-in. In the specific instance of TFAM nucleoid aggregates, these were counted using the ImageJ Particle Analysis tool (automated particle count with a minimum detected particle area threshold of 0.7 μm^2^. Parameters from TEM images (cytosol area, lysosome area, mitochondria-lysosome contact length) were measured using the ImageJ ROI manager. The number of galectolysosomes formed in mRFP-GFP-Galectin-8 cells were determined by counting GFP quenched, mRFP positive foci using the ImageJ MitoQC plug-in ^70^.

Images obtained by SoRa were 3D deconvoluted using NIS elements imaging software (Nikon, v5.21-03) and further analysed using Imaris software (v9.9.1). YFP-GAL8 foci were modelled as centroids. Similarly, the mitochondria network surface masking was applied according to the intensity of TOM20 fluorescent signal. The same modelling settings were used on all images acquired. Standard Imaris measurements were extracted showing the distance from the YFP-GAL8 foci to the rendered mitochondrial network, measured from the centre of modelled YFP-GAL8 centroids to the outer surface of the modelled mitochondrial network, with the radius of the centroid then subtracted. To assess association of YFP-GAL8 foci with the mitochondrial network, they were compared with a shuffled control. In these controls, an ImageJ plug-in was created to randomly shuffle the YFP-GAL8 foci within the cytosolic area on a per cell basis (cytosolic area as outlined by the reach of the mitochondrial network).

### Cytosol fractionation for nucleic acid quantification

Methods were adapted from previously published protocols ^47,71^. Cells were trypsinised and fresh medium added to inactivate trypsin, then centrifuged at 200 g at 4 °C. The supernatant was then aspirated, and cells washed by gentle pipetting in 1 ml of ice-cold PBS. The centrifugation and PBS wash was repeated once more. The pellet was then resuspended in 500 µl (per 60 mm dish) of ice-cold fractionation buffer (50 mM HEPES pH 7.4, 150 mM NaCl, 25 µg ml^-1^ fresh digitonin (Sigma, #11024-24-1) and 1 X Complete Protease Inhibitor Cocktail with EDTA, Roche). When RNA was to be isolated from the cytosolic fractions 4 U RNase inhibitor (Promega, #N2111) was added to the fractionation buffer. The suspensions were incubated end-over-end for 10 mins at 4 °C to allow selective plasma membrane permeabilization, then centrifuged at 1000 g for 3 min to pellet cells (insoluble fraction). The cytosolic supernatant was transferred to a fresh tube, pelleted at 1000 g for 3 mins, and the resulting cleared supernatant transferred to a fresh tube. This cytosolic supernatant was finally centrifuged at 17000 g for 10 min to pellet any remaining non-cytosolic cellular material or debris. The cytosolic fraction was transferred to a fresh tube. The insoluble fraction was solubilised in in 4% SDS lysis buffer (4% SDS, 50 mM Tris-HCl pH 7.5 and 150 mM NaCl), 300 mL of lysis buffer per 60 mm dish, and pulse sonicated.

In those instances where the cytosolic fraction was to be used for DNA abundance quantification, an additional step was introduced into the above procedure where, after initial cell trypsinisation, one-third of the cell suspension was separately pelleted and lysed by addition of 400 µl of 50 µM NaOH, followed by heating at 95°C for 30 min to solubilise DNA, and neutralised by adding 50 µl 1 M Tris-HCL (pH 8.0). These extracts served as normalisation controls (*ACTB* qPCR, see below)

### Nucleic acid purification

For total cell RNA isolation, cells were lysed directly in RLT lysis buffer with added β-mercaptoethanol as per manufacturer’s protocol (Qiagen, RNeasy kit), and RNA was isolated using by spin-column RNA extraction kit (Qiagen, RNeasy kit). For RNA isolation from cytosolic fractions (above), Trizol / chloroform extraction was performed. Briefly, cytosolic fractions were mixed with Trizol (Thermo Fisher, #15596026) in a 1:1 ratio and incubated for 5 min. 0.2 mL of chloroform per 1 mL of Trizol was added to the mixture and vortexed vigorously for 15 seconds, followed by incubation at room temperature for 2 min. The mixture was centrifuged at 12,000 g for 15 min at 4°C. Following centrifugation, the upper aqueous phase was transferred carefully into a fresh RNase-free tube. 25 μg of GlycoBlue coprecipitant (Thermo Fisher, #AM9516) was added and RNA precipitated by mixing with 0.5 mL of isopropanol per 1 ml of original Trizol reagent used, then incubation at room temperature for 10 min, followed by centrifugation at 12,000 g for 10 min at 4°C. The supernatant was removed completely and discarded. The RNA pellet was washed once with 1 ml of 75% ethanol per 1 ml of original Trizol reagent used, and subsequent centrifugation at 7,500g for 5 min at 4°C. The ethanol wash was then fully removed, and the cleaned RNA pellet air-dried for 10 min, then re-dissolved in nuclease free water. To remove trace DNA, samples were treated with DNase-I using the RNAqueous DNase kit (Thermo Fisher), according to the manufacturer’s protocol. For DNA isolation from cytosolic fractions (above), a spin-column DNA isolation kit (QIAamp, Qiagen) was used, according to the manufacturer’s protocol. DNA was eluted in nuclease-free water. Nucleic acid concentration and purity was measured using a NanoDrop 2000c spectrophotometer (Thermo Scientific).

### Quantitative PCR (qPCR)

Reverse transcription from RNA isolated from total cell lysates was done using QScript cDNA SuperMix (QuantaBio #95048), according to the manufacturer’s protocol. Reverse transcription from RNA isolated from cytosolic fractions (above) was performed using the SuperScript III reverse transcriptase (Thermo Fisher, cat.no 18080093). In both instances, random DNA hexamers were used for priming. qPCR was performed using SYBR Select Master Mix (Applied Biosystems, #4472908), according to the manufacturer’s protocol, on a StepOne Real-Time PCR System instrument (ThermoFisher). For cDNA from total cell RNA samples, transcripts were quantified using the ΔΔCt method, while standard curve methodology was used to quantify cDNA/DNA obtained from cytosolic fractions. Each independent replicate data point is typically the mean of 3 technical replicates. Primers used are listed in Table S2.

### Flow cytometry

Δ*ATG5* A549 cells were plated in 6-well plates. The following day, cells were treated with DMSO or 10 µM Pb for 24 h. 4h 20 µM Gamitrinib TPP (GTPP, MedChemExpress, HY-102007A) treatment was performed on the following day prior to incubation with JC-10. After corresponding times, cells were incubated with 15 µM JC-10 (Enzo, #ENZ-52305) or 50 nM LysoTracker™ Red DND-99 (Invitrogen, #L7528) in non-phenol red DMEM media (Gibco, #31053028) (supplemented with 10% FBS, 2 mM L-glutamine and 10 U ml^-1^ 10 μg ml^-1^ penicillin/streptomycin) for 30 min at 37°C. Cells were then washed with PBS, trypsinised for 5 min and fresh flow cytometry buffer (non-phenol red DMEM media supplemented with 1% FBS, 2 mM L-glutamine and 10 U ml^-1^ 10 μg ml^-1^ penicillin/streptomycin) was added to stop the reaction. Cells were centrifuged at 150 g for 3 min, resuspended in 500 μl of flow cytometry buffer and kept on ice until analysed in a Cytoflex S flow cytometer (Beckman Coulter). JC-10 monomeric was excited by a 488 nm laser line and emission collected through a 525/40BP. JC-aggregates were excited with a 561 nm laser line and emission was collected through 585/42BP. To quantify the changes in mitochondrial membrane potential the fluorescent ratio of aggregating/monomeric JC-10 was used. Lysotracker red was excited by a 561 nm laser line and its emission was collected through 610/20BP. Data was exported and analysed using FlowJo v10.10 software.

### Statistical analyses

All statistical analysis was carried out using GraphPad (Prism) (version 9.5.1). Sample size for experiments to provide adequate power were predicted based upon prior experience of the experimenter and/or pilot data sampling. All independent biological replicates are represented with standard error of the mean (s.e.m.). In the case of electron microscopy image analyses, individual cells are represented as data points and variance described as standard deviation (s.d.). Tests are detailed in figure legends. In general, an assumption of normality was made and all tests were two-sided. Thus, where pairwise comparisons between two samples are performed, this is performed with an unpaired Student’s t-test or a one-sample t-test (latter instance when comparing against the normalisation control). In a minority of cases where a non-normal distribution was clearly observed, a one-sample Wilcoxon test was instead used for pairwise comparison (as indicated in figure legends). When multiple comparisons were performed in instances where there was only one independent variable, we generally used one-way ANOVA with Holm-Šídák testing. Where multiple comparisons were performed in instances of two independent variables in data set (e.g., cell line identity and/or drug application) we generally performed two-way ANOVA with Holm-Šídák testing. In the case of cytosolic RNA/DNA analysis, ratio paired t-tests were performed. Replicates containing outliers as detected by the Grubb’s test were removed prior to the ratio paired t-test. Comparison of pairs of frequency distributions were performed by the Kolmogorov-Smirnov test for each of three replicates, and a *P*-value for each replicate generated individually.

## Code availability

The custom FIJI (ImageJ) plug-in macro to shuffle galectin-8 foci within cell space is reproduced below:

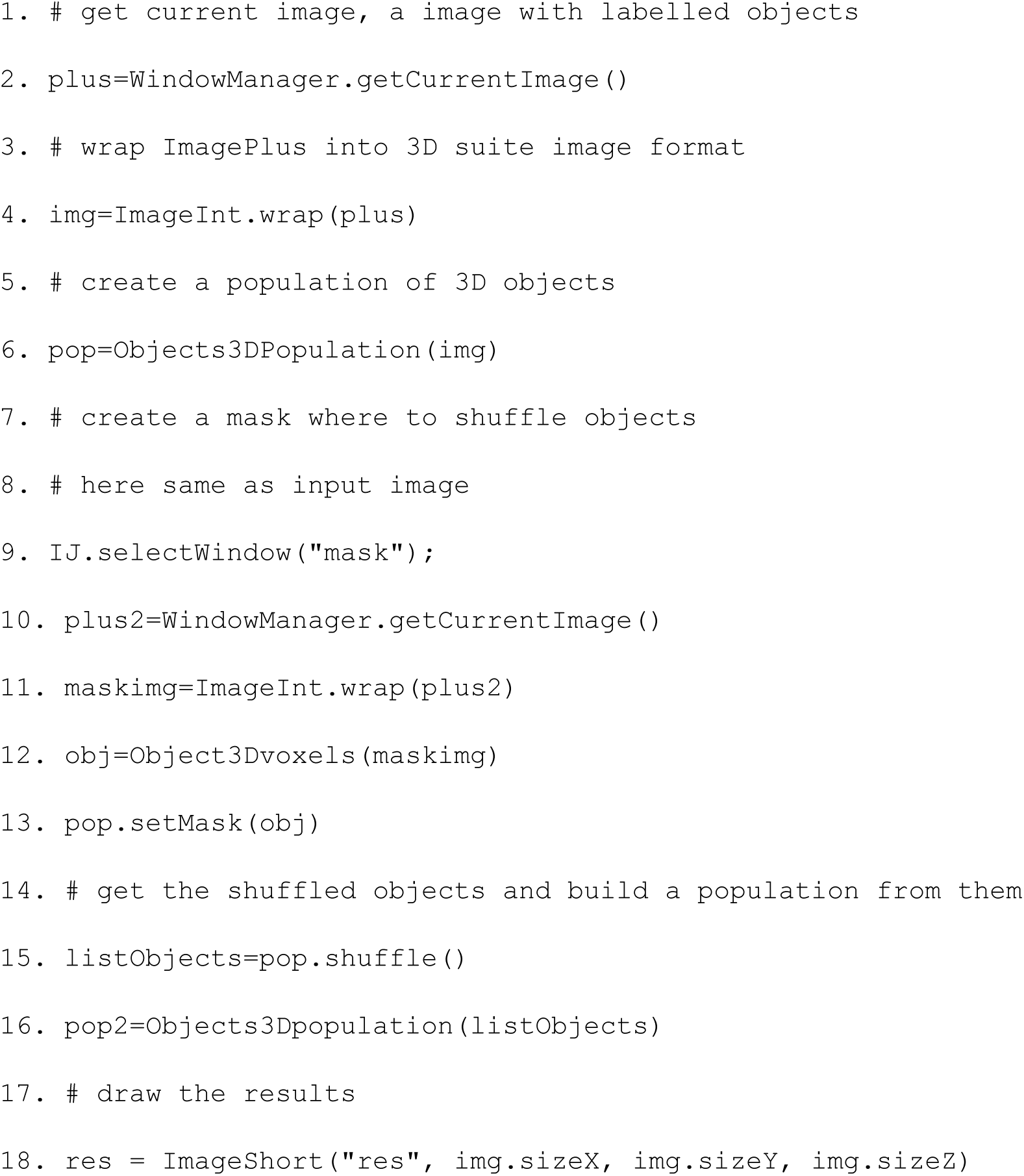
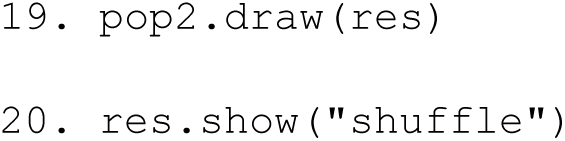

## Data Availability

All relevant data and details of resources can be found within the article and its supplementary information. Uncropped blots are shown in Figs. S4-6. Raw data not included in this manuscript are available from the corresponding author upon reasonable request. Plasmids, cell lines and other reagents are available from the corresponding author upon reasonable request.

## Additional Information

The author(s) declare no competing interests.

## Author contributions

Study conception and funding acquisition by SW. Experiments designed by MB and SW. Experiments performed by MB, with additional data generated by CSC, AK, AJK, TEL, KWI and NJM. Refinement of mitochondrial nucleic acid analysis and mitochondrial genome depletion experiments by AD. FIJI macro for GAL8-mitochondrion image analysis written by LM. Time-lapse video microscopy by AW. All data analysis by MB, TEL, CSC and AK. Manuscript drafted by MB and SW. Editing performed by MB, NJM and SW.

## Notes

### Competing Interest Statement

The authors have declared no competing interest.

### Summary of Updates

Figure 1 separated into two figures for clarity. New data: Lysotracker Red and JC-10 flow cytometry analysis of the effects of Palbociclib on lysosomes and mitochondrial membrane potential. Some minor text edits for clarity. New authors added (CSC and AK)

## References

1 Zoncu, R. & Perera, R. M. Built to last: lysosome remodeling and repair in health and disease. Trends Cell Biol 32, 597–610, doi:10.1016/j.tcb.2021.12.009 (2022).

2 Thurston, T. L., Wandel, M. P., von Muhlinen, N., Foeglein, A. & Randow, F. Galectin 8 targets damaged vesicles for autophagy to defend cells against bacterial invasion. Nature 482, 414–418, doi:10.1038/nature10744 (2012).

3 Wang, F., Gomez-Sintes, R. & Boya, P. Lysosomal membrane permeabilization and cell death. Traffic 19, 918–931, doi:10.1111/tra.12613 (2018).

4 Radulovic, M. et al. ESCRT-mediated lysosome repair precedes lysophagy and promotes cell survival. EMBO J 37, e99753, doi:10.15252/embj.201899753 (2018).

5 Skowyra, M. L., Schlesinger, P. H., Naismith, T. V. & Hanson, P. I. Triggered recruitment of ESCRT machinery promotes endolysosomal repair. Science 360, doi:10.1126/science.aar5078 (2018).

6 Tan, J. X. & Finkel, T. A phosphoinositide signalling pathway mediates rapid lysosomal repair. Nature 609, 815–821, doi:10.1038/s41586-022-05164-4 (2022).

7 Jia, J. et al. Galectin-3 Coordinates a Cellular System for Lysosomal Repair and Removal. Dev Cell 52, 69–87 e68, doi:10.1016/j.devcel.2019.10.025 (2020).

8 Papadopoulos, C., Kravic, B. & Meyer, H. Repair or Lysophagy: Dealing with Damaged Lysosomes. J Mol Biol 432, 231–239, doi:10.1016/j.jmb.2019.08.010 (2020).

9 Maejima, I. et al. Autophagy sequesters damaged lysosomes to control lysosomal biogenesis and kidney injury. EMBO J 32, 2336–2347, doi:10.1038/emboj.2013.171 (2013).

10 Dikic, I. & Elazar, Z. Mechanism and medical implications of mammalian autophagy. Nat Rev Mol Cell Biol 19, 349–364, doi:10.1038/s41580-018-0003-4 (2018).

11 Vargas, J. N. S., Hamasaki, M., Kawabata, T., Youle, R. J. & Yoshimori, T. The mechanisms and roles of selective autophagy in mammals. Nat Rev Mol Cell Biol 24, 167–185, doi:10.1038/s41580-022-00542-2 (2023).

12 Hoyer, M. J., Swarup, S. & Harper, J. W. Mechanisms Controlling Selective Elimination of Damaged Lysosomes. Curr Opin Physiol 29, 100590, doi:10.1016/j.cophys.2022.100590 (2022).

13 Jia, J. et al. Galectins Control mTOR in Response to Endomembrane Damage. Mol Cell 70, 120–135 e128, doi:10.1016/j.molcel.2018.03.009 (2018).

14 Gahlot, P. et al. Lysosomal damage sensing and lysophagy initiation by SPG20-ITCH. Mol Cell 84, 1556–1569 e1510, doi:10.1016/j.molcel.2024.02.029 (2024).

15 Kravic, B. et al. Ubiquitin profiling of lysophagy identifies actin stabilizer CNN2 as a target of VCP/p97 and uncovers a link to HSPB1. Mol Cell 82, 2633–2649 e2637, doi:10.1016/j.molcel.2022.06.012 (2022).

16 Papadopoulos, C. et al. VCP/p97 cooperates with YOD1, UBXD1 and PLAA to drive clearance of ruptured lysosomes by autophagy. EMBO J 36, 135–150, doi:10.15252/embj.201695148 (2017).

17 Eapen, V. V., Swarup, S., Hoyer, M. J., Paulo, J. A. & Harper, J. W. Quantitative proteomics reveals the selectivity of ubiquitin-binding autophagy receptors in the turnover of damaged lysosomes by lysophagy. Elife 10, e72328, doi:10.7554/eLife.72328 (2021).

18 Gallagher, E. R. & Holzbaur, E. L. F. The selective autophagy adaptor p62/SQSTM1 forms phase condensates regulated by HSP27 that facilitate the clearance of damaged lysosomes via lysophagy. Cell Rep 42, 112037, doi:10.1016/j.celrep.2023.112037 (2023).

19 Chauhan, S. et al. TRIMs and Galectins Globally Cooperate and TRIM16 and Galectin-3 Co-direct Autophagy in Endomembrane Damage Homeostasis. Dev Cell 39, 13–27, doi:10.1016/j.devcel.2016.08.003 (2016).

20 Jia, J. et al. AMPK, a Regulator of Metabolism and Autophagy, Is Activated by Lysosomal Damage via a Novel Galectin-Directed Ubiquitin Signal Transduction System. Mol Cell 77, 951–969 e959, doi:10.1016/j.molcel.2019.12.028 (2020).

21 Park, N. Y. et al. Activation of lysophagy by a TBK1-SCF(FBXO3)-TMEM192-TAX1BP1 axis in response to lysosomal damage. Nat Commun 16, 1109, doi:10.1038/s41467-025-56294-y (2025).

22 Johansen, T. & Lamark, T. Selective Autophagy: ATG8 Family Proteins, LIR Motifs and Cargo Receptors. J Mol Biol 432, 80–103, doi:10.1016/j.jmb.2019.07.016 (2020).

23 Lazarou, M. et al. The ubiquitin kinase PINK1 recruits autophagy receptors to induce mitophagy. Nature 524, 309–314, doi:10.1038/nature14893 (2015).

24 Vargas, J. N. S. et al. Spatiotemporal Control of ULK1 Activation by NDP52 and TBK1 during Selective Autophagy. Mol Cell 74, 347–362 e346, doi:10.1016/j.molcel.2019.02.010 (2019).

25 Turco, E. et al. FIP200 Claw Domain Binding to p62 Promotes Autophagosome Formation at Ubiquitin Condensates. Mol Cell 74, 330–346 e311, doi:10.1016/j.molcel.2019.01.035 (2019).

26 Ravenhill, B. J. et al. The Cargo Receptor NDP52 Initiates Selective Autophagy by Recruiting the ULK Complex to Cytosol-Invading Bacteria. Mol Cell 74, 320–329 e326, doi:10.1016/j.molcel.2019.01.041 (2019).

27 Bhattacharya, A. et al. A lysosome membrane regeneration pathway depends on TBC1D15 and autophagic lysosomal reformation proteins. Nat Cell Biol 25, 685–698, doi:10.1038/s41556-023-01125-9 (2023).

28 Cross, J. et al. Lysosome damage triggers direct ATG8 conjugation and ATG2 engagement via non-canonical autophagy. J Cell Biol 222, doi:10.1083/jcb.202303078 (2023).

29 Eriksson, I., Waster, P. & Ollinger, K. Restoration of lysosomal function after damage is accompanied by recycling of lysosomal membrane proteins. Cell Death Dis 11, 370, doi:10.1038/s41419-020-2527-8 (2020).

30 Jia, J. et al. Stress granules and mTOR are regulated by membrane atg8ylation during lysosomal damage. J Cell Biol 221, doi:10.1083/jcb.202207091 (2022).

31 Duran, J. et al. Calcium signaling from damaged lysosomes induces cytoprotective stress granules. EMBO J 43, 6410–6443, doi:10.1038/s44318-024-00292-1 (2024).

32 Neuwirt, E. et al. Tyrosine kinase inhibitors can activate the NLRP3 inflammasome in myeloid cells through lysosomal damage and cell lysis. Sci Signal 16, eabh1083, doi:10.1126/scisignal.abh1083 (2023).

33 Llanos, S. et al. Lysosomal trapping of palbociclib and its functional implications. Oncogene 38, 3886–3902, doi:10.1038/s41388-019-0695-8 (2019).

34 Goel, S. et al. CDK4/6 inhibition triggers anti-tumour immunity. Nature 548, 471–475, doi:10.1038/nature23465 (2017).

35 Fan, H. et al. DNA damage induced by CDK4 and CDK6 blockade triggers anti-tumor immune responses through cGAS-STING pathway. Commun Biol 6, 1041, doi:10.1038/s42003-023-05412-x (2023).

36 Wu, J. et al. Separable Cell Cycle Arrest and Immune Response Elicited through Pharmacological CDK4/6 and MEK Inhibition in RASmut Disease Models. Mol Cancer Ther 23, 1801–1814, doi:10.1158/1535-7163.MCT-24-0369 (2024).

37 Roulois, D. et al. DNA-Demethylating Agents Target Colorectal Cancer Cells by Inducing Viral Mimicry by Endogenous Transcripts. Cell 162, 961–973, doi:10.1016/j.cell.2015.07.056 (2015).

38 Akira, S., Uematsu, S. & Takeuchi, O. Pathogen recognition and innate immunity. Cell 124, 783–801, doi:10.1016/j.cell.2006.02.015 (2006).

39 Yoneyama, M. et al. The RNA helicase RIG-I has an essential function in double-stranded RNA-induced innate antiviral responses. Nat Immunol 5, 730–737, doi:10.1038/ni1087 (2004).

40 Seth, R. B., Sun, L., Ea, C. K. & Chen, Z. J. Identification and characterization of MAVS, a mitochondrial antiviral signaling protein that activates NF-kappaB and IRF 3. Cell 122, 669–682, doi:10.1016/j.cell.2005.08.012 (2005).

41 Rehwinkel, J. & Gack, M. U. RIG-I-like receptors: their regulation and roles in RNA sensing. Nat Rev Immunol 20, 537–551, doi:10.1038/s41577-020-0288-3 (2020).

42 Sun, L., Wu, J., Du, F., Chen, X. & Chen, Z. J. Cyclic GMP-AMP synthase is a cytosolic DNA sensor that activates the type I interferon pathway. Science 339, 786–791, doi:10.1126/science.1232458 (2013).

43 Crow, Y. J. & Stetson, D. B. The type I interferonopathies: 10 years on. Nat Rev Immunol 22, 471–483, doi:10.1038/s41577-021-00633-9 (2022).

44 Zitvogel, L., Galluzzi, L., Kepp, O., Smyth, M. J. & Kroemer, G. Type I interferons in anticancer immunity. Nat Rev Immunol 15, 405–414, doi:10.1038/nri3845 (2015).

45 Ishikawa, H. & Barber, G. N. STING is an endoplasmic reticulum adaptor that facilitates innate immune signalling. Nature 455, 674–678, doi:10.1038/nature07317 (2008).

46 Mackenzie, K. J. et al. cGAS surveillance of micronuclei links genome instability to innate immunity. Nature 548, 461–465, doi:10.1038/nature23449 (2017).

47 Dhir, A. et al. Mitochondrial double-stranded RNA triggers antiviral signalling in humans. Nature 560, 238–242, doi:10.1038/s41586-018-0363-0 (2018).

48 Tigano, M., Vargas, D. C., Tremblay-Belzile, S., Fu, Y. & Sfeir, A. Nuclear sensing of breaks in mitochondrial DNA enhances immune surveillance. Nature 591, 477–481, doi:10.1038/s41586-021-03269-w (2021).

49 Katayama, H. et al. Visualizing and Modulating Mitophagy for Therapeutic Studies of Neurodegeneration. Cell 181, 1176–1187 e1116, doi:10.1016/j.cell.2020.04.025 (2020).

50 Jimenez-Moreno, N., Salomo-Coll, C., Murphy, L. C. & Wilkinson, S. Signal-Retaining Autophagy Indicator as a Quantitative Imaging Method for ER-Phagy. Cells 12, doi:10.3390/cells12081134 (2023).

51 Wong, Y. C., Ysselstein, D. & Krainc, D. Mitochondria-lysosome contacts regulate mitochondrial fission via RAB7 GTP hydrolysis. Nature 554, 382–386, doi:10.1038/nature25486 (2018).

52 Uzhachenko, R. V. et al. Metabolic modulation by CDK4/6 inhibitor promotes chemokine-mediated recruitment of T cells into mammary tumors. Cell Rep 35, 109271, doi:10.1016/j.celrep.2021.109271 (2021).

53 West, A. P. et al. Mitochondrial DNA stress primes the antiviral innate immune response. Nature 520, 553–557, doi:10.1038/nature14156 (2015).

54 Kim, J. et al. VDAC oligomers form mitochondrial pores to release mtDNA fragments and promote lupus-like disease. Science 366, 1531–1536, doi:10.1126/science.aav4011 (2019).

55 Torres-Odio, S. et al. Loss of Mitochondrial Protease CLPP Activates Type I IFN Responses through the Mitochondrial DNA-cGAS-STING Signaling Axis. J Immunol 206, 1890–1900, doi:10.4049/jimmunol.2001016 (2021).

56 Antonicka, H., Sasarman, F., Nishimura, T., Paupe, V. & Shoubridge, E. A. The mitochondrial RNA-binding protein GRSF1 localizes to RNA granules and is required for posttranscriptional mitochondrial gene expression. Cell Metab 17, 386–398, doi:10.1016/j.cmet.2013.02.006 (2013).

57 Newman, A. C., Kemp, A. J., Drabsch, Y., Behrends, C. & Wilkinson, S. Autophagy acts through TRAF3 and RELB to regulate gene expression via antagonism of SMAD proteins. Nat Commun 8, 1537, doi:10.1038/s41467-017-00859-z (2017).

58 Riley, J. S. et al. Mitochondrial inner membrane permeabilisation enables mtDNA release during apoptosis. EMBO J 37, doi:10.15252/embj.201899238 (2018).

59 McArthur, K. et al. BAK/BAX macropores facilitate mitochondrial herniation and mtDNA efflux during apoptosis. Science 359, doi:10.1126/science.aao6047 (2018).

60 Kang, B. H. et al. Combinatorial drug design targeting multiple cancer signaling networks controlled by mitochondrial Hsp90. J Clin Invest 119, 454–464, doi:10.1172/JCI37613 (2009).

61 Voeltz, G. K., Sawyer, E. M., Hajnoczky, G. & Prinz, W. A. Making the connection: How membrane contact sites have changed our view of organelle biology. Cell 187, 257–270, doi:10.1016/j.cell.2023.11.040 (2024).

62 Xu, L. et al. Salmonella Induces the cGAS-STING-Dependent Type I Interferon Response in Murine Macrophages by Triggering mtDNA Release. mBio 13, e0363221, doi:10.1128/mbio.03632-21 (2022).

63 Bussi, C. et al. Lysosomal damage drives mitochondrial proteome remodelling and reprograms macrophage immunometabolism. Nat Commun 13, 7338, doi:10.1038/s41467-022-34632-8 (2022).

64 Nakahira, K. et al. Autophagy proteins regulate innate immune responses by inhibiting the release of mitochondrial DNA mediated by the NALP3 inflammasome. Nat Immunol 12, 222–230, doi:10.1038/ni.1980 (2011).

65 Mathew, R. et al. Functional role of autophagy-mediated proteome remodeling in cell survival signaling and innate immunity. Mol Cell 55, 916–930, doi:10.1016/j.molcel.2014.07.019 (2014).

66 Prabakaran, T. et al. Attenuation of cGAS-STING signaling is mediated by a p62/SQSTM1-dependent autophagy pathway activated by TBK1. EMBO J 37, doi:10.15252/embj.201797858 (2018).

67 He, X. et al. RNF34 functions in immunity and selective mitophagy by targeting MAVS for autophagic degradation. EMBO J 38, e100978, doi:10.15252/embj.2018100978 (2019).

68 Jimenez-Loygorri, J. I. et al. Mitophagy curtails cytosolic mtDNA-dependent activation of cGAS/STING inflammation during aging. Nat Commun 15, 830, doi:10.1038/s41467-024-45044-1 (2024).

69 Newman, A. C. et al. TBK1 kinase addiction in lung cancer cells is mediated via autophagy of Tax1bp1/Ndp52 and non-canonical NF-kappaB signalling. PLoS One 7, e50672, doi:10.1371/journal.pone.0050672 (2012).

70 Montava-Garriga, L., Singh, F., Ball, G. & Ganley, I. G. Semi-automated quantitation of mitophagy in cells and tissues. Mech Ageing Dev 185, 111196, doi:10.1016/j.mad.2019.111196 (2020).

71 Bryant, J. D., Lei, Y., VanPortfliet, J. J., Winters, A. D. & West, A. P. Assessing Mitochondrial DNA Release into the Cytosol and Subsequent Activation of Innate Immune-related Pathways in Mammalian Cells. Curr Protoc 2, e372, doi:10.1002/cpz1.372 (2022).

72 Ran, F. A. et al. Genome engineering using the CRISPR-Cas9 system. Nat Protoc 8, 2281–2308, doi:10.1038/nprot.2013.143 (2013).

73 Yang, X. et al. A public genome-scale lentiviral expression library of human ORFs. Nat Methods 8, 659–661, doi:10.1038/nmeth.1638 (2011).

74 Campeau, E. et al. A versatile viral system for expression and depletion of proteins in mammalian cells. PLoS One 4, e6529, doi:10.1371/journal.pone.0006529 (2009).

