## Supplemental Figures and Legends for "Chronic lysosome damage boosts interferon responses to Palbociclib dependent upon the mitochondrion"

### Supplementary Materials

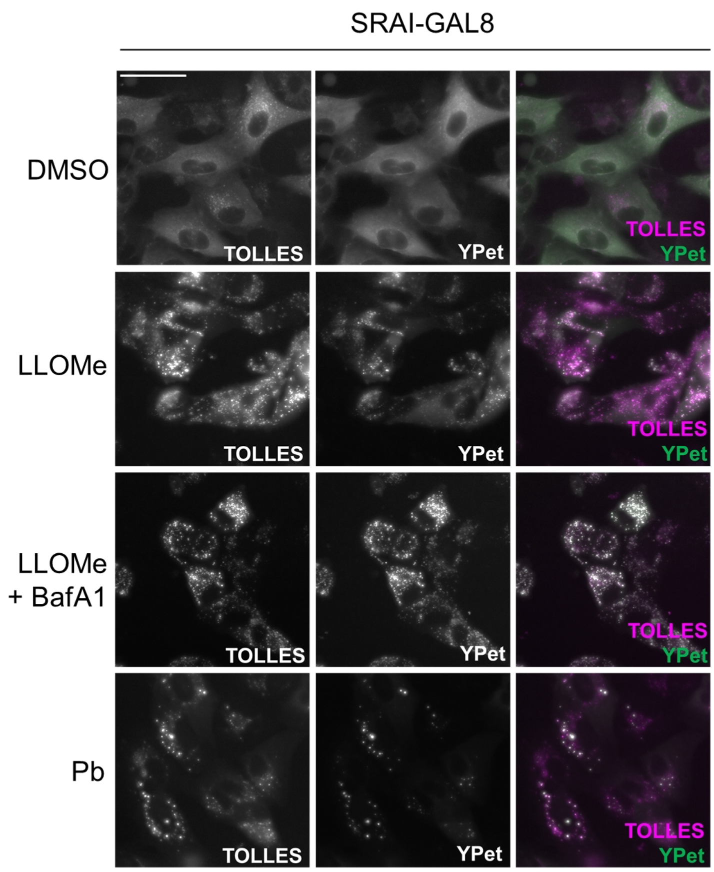


**Figure S1**

A549- or MCF7-SRAI-GAL8 cells were treated for 24 h with DMSO, 0.7 mM LLOMe (+/- 0.2 μM of the lysosome blocker Bafilomycin, BafA1), or 10 μM Palbociclib, Pb, then imaged by widefield fluorescence microscopy. TOLLES-only foci are generated in response to both drugs, but this is blocked by BafA1 (representative of n = 3 independent replicates).

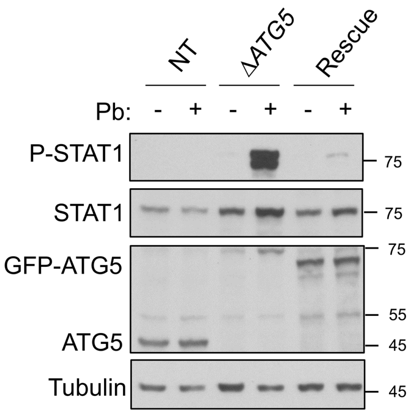


**Figure S2**

A549 NT (non-targeted wild-type), Δ*ATG5* or Rescue cells (Δ*ATG5* + *GFP-ATG5*) were treated with DMSO (-) or 10 μM Palbociclib, Pb (+), for 48 h, then immunoblotted to assess interferon pathway activation (phospho-Y701 STAT1, P-STAT1) (n = 3, representative replicate shown).

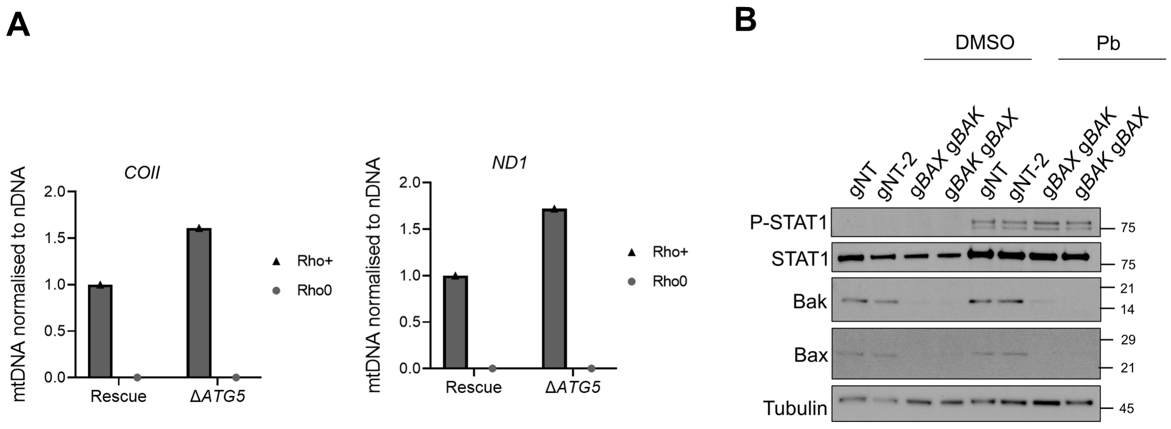


**Figure S3**

**A)** Rho+ and Rho0 cultures of A549 Rescue (Δ*ATG5* + GFP-*ATG5*) and Δ*ATG5* cells generated as described in **Fig. 7A, B, C** were subjected to total cellular DNA isolation and qPCR for mitochondrial DNA (mtDNA) sequences *COII* and *ND1* (n = 1, normalised to *ACTB*, nuclear DNA, nDNA).

**B)** A549 Δ*ATG5* cells stably-expressing Cas9 were transduced with non-targeting sgRNA (gNT or gNT-2) or sequentially transduced with gRNAs against *BAX* and *BAK* in denoted order. Cells were treated with DMSO or 10 μM Pb for 48 h and immunoblotted for phospho-Y701-STAT1 (P-STAT1) and Bax/Bak knockout controls (representative of n = 3 independent replicates).

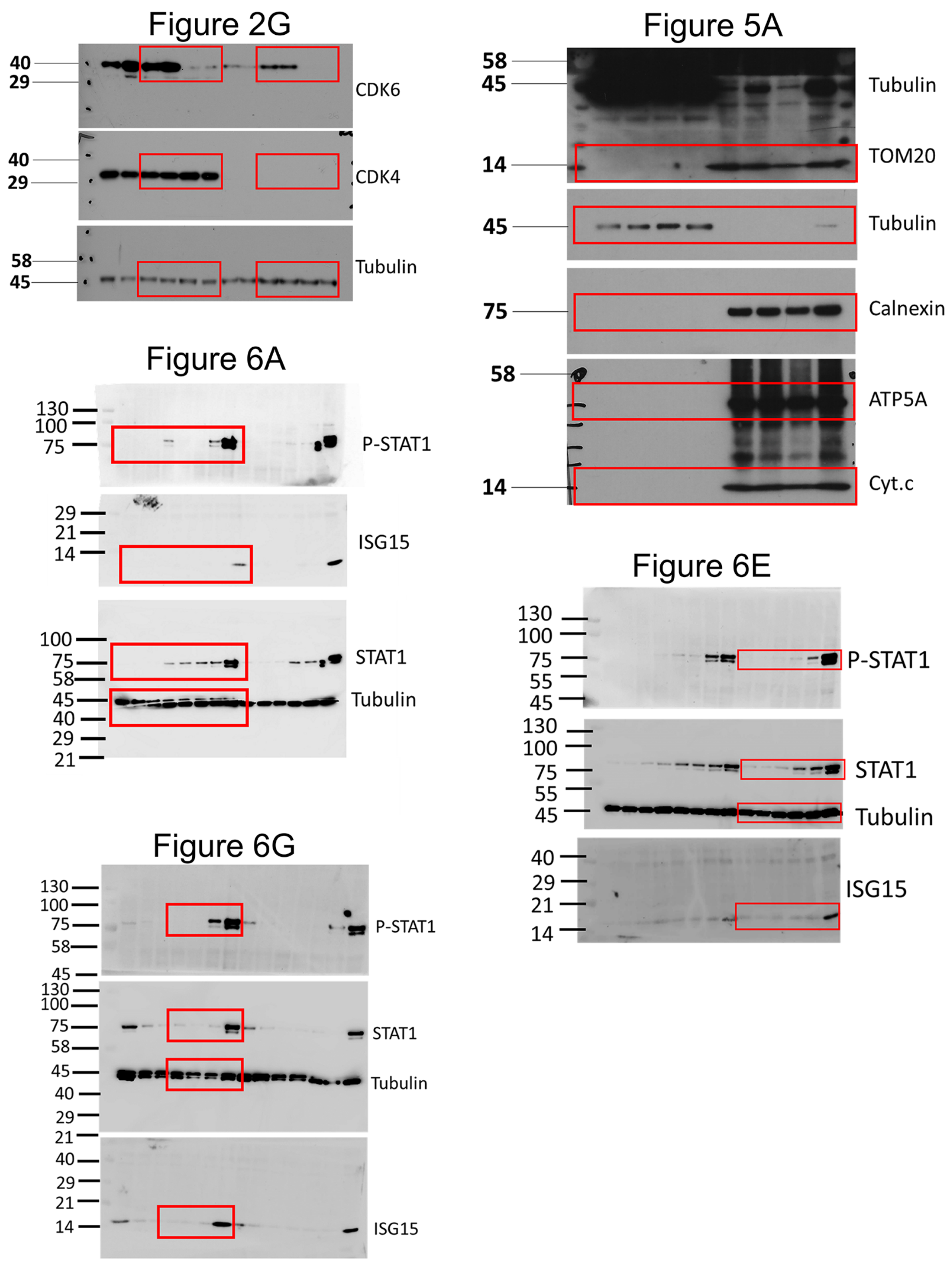


**Figure S4**

Raw blot data, 1 of 3, see also Figures S5 and S6.

Figure 6D

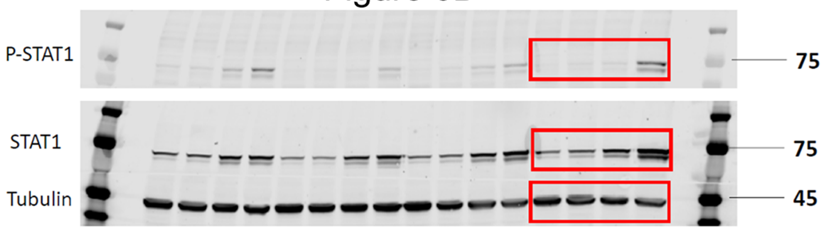


Figure 6I

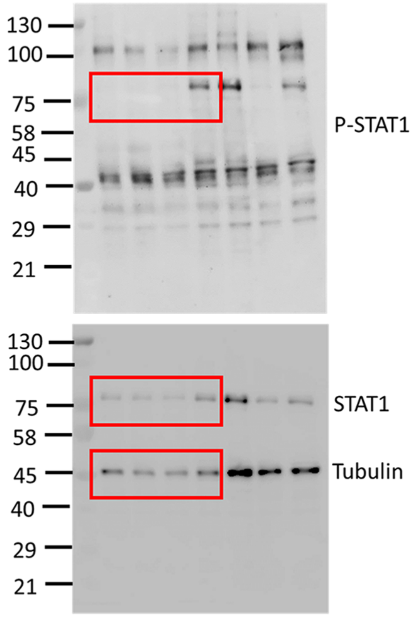


Figure 7A

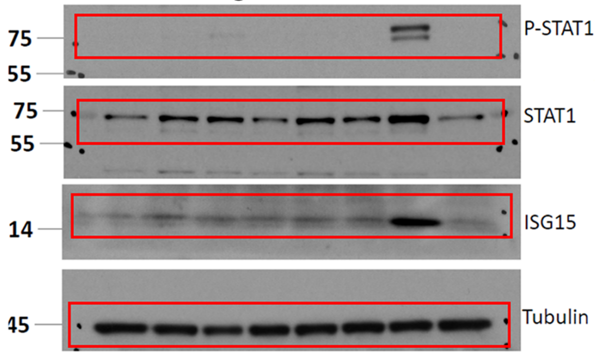


Figure 7D

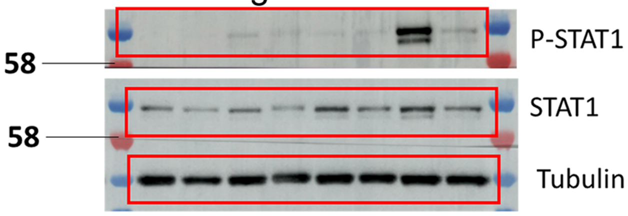


Figure 7F

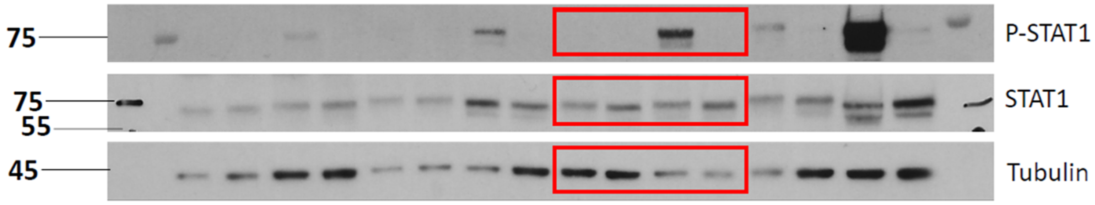


**Figure S5**

Raw blot data, 2 of 3, see also Figures S4 and S6.

Figure 7I

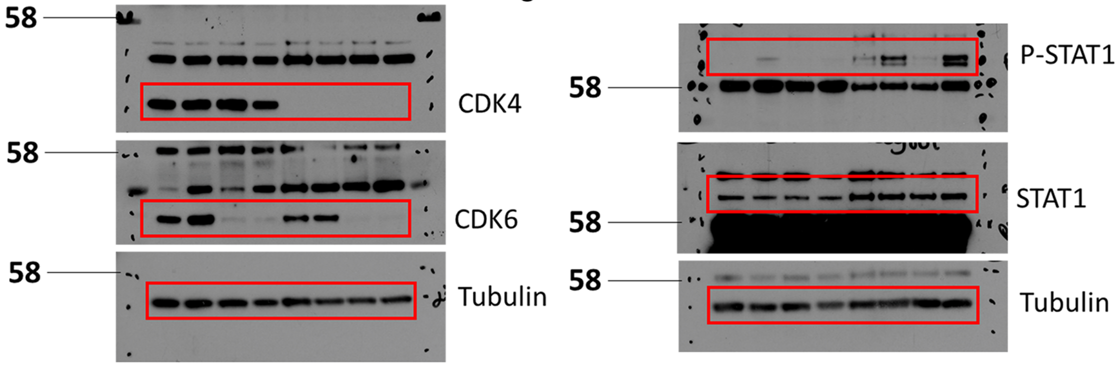


Supp Fig 2

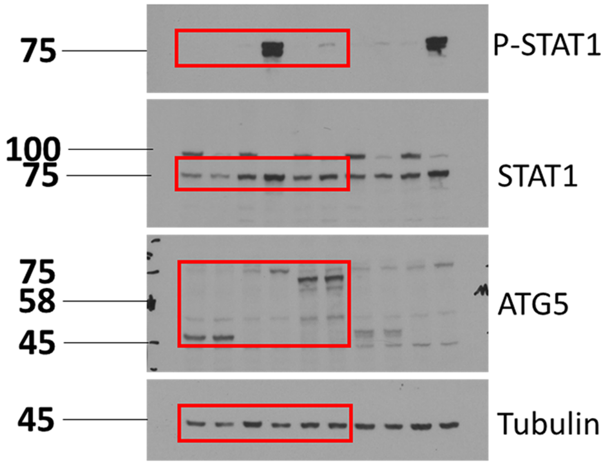


Supp Fig 3B

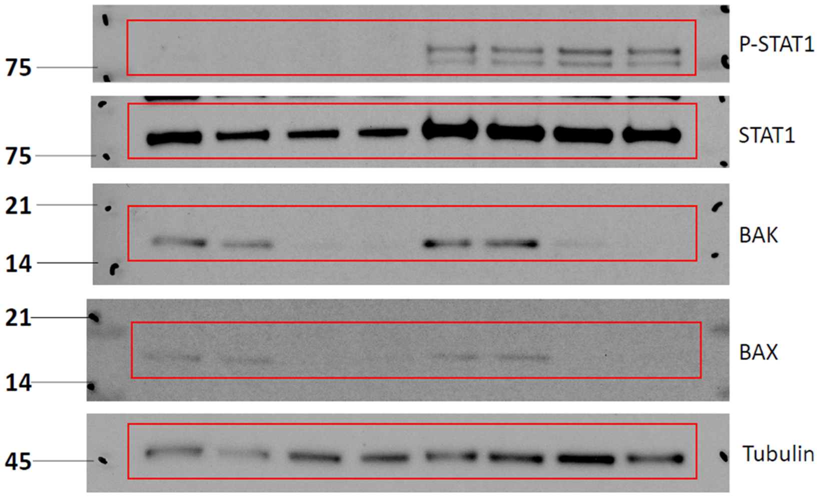


**Figure S6**

Raw blot data, 3 of 3, see also Figures S4 and S5.

### Supplementary Legends

#### Movie S1

180° rotation around y-axis of 3D rendered cell shown in **Fig. 3E** (equivalent to lower zoom panel 1). YFP-GAL8 in green, TOM20 in red. Mitochondrial volume has been rendered to model the space occupied by the mitochondrial network.

#### Movie S2

Corresponds to still frames in **Fig. 3G** row 1. YFP-GAL8 in green, MitoTracker in red.

#### Movie S3

Corresponds to still frames in **Fig. 3G** row 2. YFP-GAL8 in green, MitoTracker in red.

#### Table S1

Details of plasmids used in this study.

#### Table S2

Details of oligonucleotides used in this study.
